# A class of antibodies that overcome a steric barrier to cross-group neutralization of influenza viruses

**DOI:** 10.1101/2023.04.01.535242

**Authors:** Holly C. Simmons, Akiko Watanabe, Thomas H. Oguin, Gregory D. Sempowski, Masayuki Kuraoka, Garnett Kelsoe, Kevin R. McCarthy

## Abstract

Antibody titers that inhibit the influenza virus hemagglutinin (HA) from engaging its receptor are the accepted correlate of protection from infection. Many potent antibodies with broad, intra-subtype specificity bind HA at the receptor binding site (RBS). One barrier to broad H1-H3 cross-subtype neutralization is an insertion (133a) between positions 133 and 134 on the rim of the H1 HA RBS. We describe here a class of antibodies that overcome this barrier. These genetically unrestricted antibodies are abundant in the human B-cell memory compartment. Analysis of the affinities of selected members of this class for historical H1 and H3 isolates suggest that they were elicited by H3 exposure and broadened or diverted by later exposure(s) to H1 HA. RBS mutations in egg-adapted vaccine strains cause the new H1 specificity of these antibodies to depend on the egg adaptation. The results suggest that suitable immunogens might elicit 133a-independent, H1-H3 cross neutralization by RBS-directed antibodies.

## Introduction

Influenza virus hemagglutinin protein (HA) mediates cellular attachment by engaging sialic acid adducts on glycoproteins and glycolipids. The titer of circulating antibodies that inhibit this interaction correlates with protection from infection^1^. Immunity to influenza viruses is limited by large antigenic divisions between the 18 influenza A HA serotypes, which are classified as either group 1 (H1, 2, 5, 6, 8, 9, 11, 12, 13, 16, 17, 18) or group 2 (H3, 4, 7, 10, 14, 15). As a result, humans do not mount broadly protective responses to influenza. Moreover, continuous antigenic evolution erodes immunity elicited by prior strains of any specific serotype. The strain composition of vaccines to human H1 and H3 viruses must therefore be reformulated on a nearly annual basis. Broadly protective antibodies that engage conserved epitopes on HA can nonetheless be identified in human repertoires^2^, and immunogens that selectively elicit these antibodies might confer broader, and longer lasting immunity than do current vaccines.

One group of broadly neutralizing influenza antibodies engage the HA receptor binding site (RBS) with sialic acid-like contacts^2-7^. RBS-directed antibodies that neutralize decades of antigenic variation within a single serotype appear to be more common than those that cross-neutralize H1N1 and H3N2 viruses^2-10^. An insertion at the RBS periphery in H1 HAs (at a position designated 133a to adhere to H3 amino acid numbering) creates a steric block to H1-H3 neutralization by RBS-directed antibodies^5,8,10-12^. The insertion was lost between 1995 and 2009, and cross-neutralizing antibodies have been described that engage H1 HAs that circulated during this time interval^5,8,9^. The 2009 H1N1 pandemic reintroduced an HA with 133a, however, and potent H1-H3 cross-neutralization with this virus or its descendants has not been reported.

The RBS coordinates the terminal sialic acid of a glycan chain. Human influenza viruses have a strong preference for α2,6-linked sialic acids, while avian viruses prefer α2,3-linkages^13,14^. Propagation of human viruses in chicken eggs, the major source of vaccine material, selects for mutants that more efficiently engage α2,3 receptors. These mutations occur within the RBS and can impact RBS-directed antibody binding^15-19^. One common substitution alters the conformation of the RBS^16^, such that immunization with egg-adapted HAs may elicit immunity specific for the vaccine component, and the resulting antibodies then fail to engage the circulating virus^15-18,20,21^. These mutations limit vaccine efficacy and specifically interfere with the development of RBS-directed antibody responses.

We define here. a previously unrecognized class of RBS-directed antibodies that are abundant in circulating human memory B cells (Bmem). Members of this class have a common motif in HCDR3 that mimics many of the authentic receptor contacts. The antibodies we examined neutralized certain H3 and H1 strains, including some H1 HAs with the K133a insertion and some without. Thus, they illustrate that the human immune system can surmount the steric barrier to cross-neutralization generated by the 133a insertion. Their widespread distribution suggests that the barrier may be relatively low. For the two antibodies we characterized in detail, from donors of different ages and different geographic locations, reactivity with historical H1 isolates appears to have extended from the early years of the 21st century through about 2015; the reactivity with historical H3 isolates spanned the late 1980s to the late 1990s. For one of the antibodies, H1 binding and neutralization depended on the mutation Q226R, which was present only in the vaccine strains as a result of adaptation to growth in eggs. These data suggest that an H3-specific B cell can affinity mature, upon exposure to H1 by infection or vaccination, to recognize an H1 HA and that the H1 adaptation need not depend on the presence or absence of a K133a insertion.

## Results

### K03.28

We earlier we profiled the HA reactivities of hundreds of human antibodies, secreted from single circulating Bmem cells^8,22^. We determined that cross-serotype and cross-group binding antibodies were abundant. Antibody competition assays showed that a particular antibody example, K03.28, from donor KEL03, bound HA at the RBS; it bound H1 and H3 HAs, and bound H1 HAs with and without the 133a insertion^8^. We have now confirmed that K03.28 potently neutralizes H1N1 and H3N2 viruses (Figure S1).

We have now determined the structure of K03.28 bound with the HA head domain of A/California/07/2009 (H1N1)(NYMC-X181), (H1 X-181), (Figure 1A). Its 19-residue HCDR3 projects into the RBS, making extensive contacts with conserved sialic acid coordinating residues. In particular, an amino acid triplet at the tip of HCDR3, 107-Glu-Gly-Trp-109, mimics contacts made by the sialic acid receptor (Figure 1A). Polar interactions between sialic acid hydroxyls 7 and 8 with HA Tyr-98 and His-183 are approximated by the carboxyl group of Glu-107. Glycine 108 does not directly contact HA, but its positive phi angle orients Trp-109 to create a pi interaction with Trp-153 of HA and to emulate contacts made by the acetamido group of sialic acid. These include van der Waals contacts with HA residues Trp-153, Leu-194 and Val-155 and a hydrogen bond from the ε1 nitrogen to the carbonyl of Val-135. To our knowledge, receptor mimicry by this tripartite motif has not previously been described.

**Figure 1.**
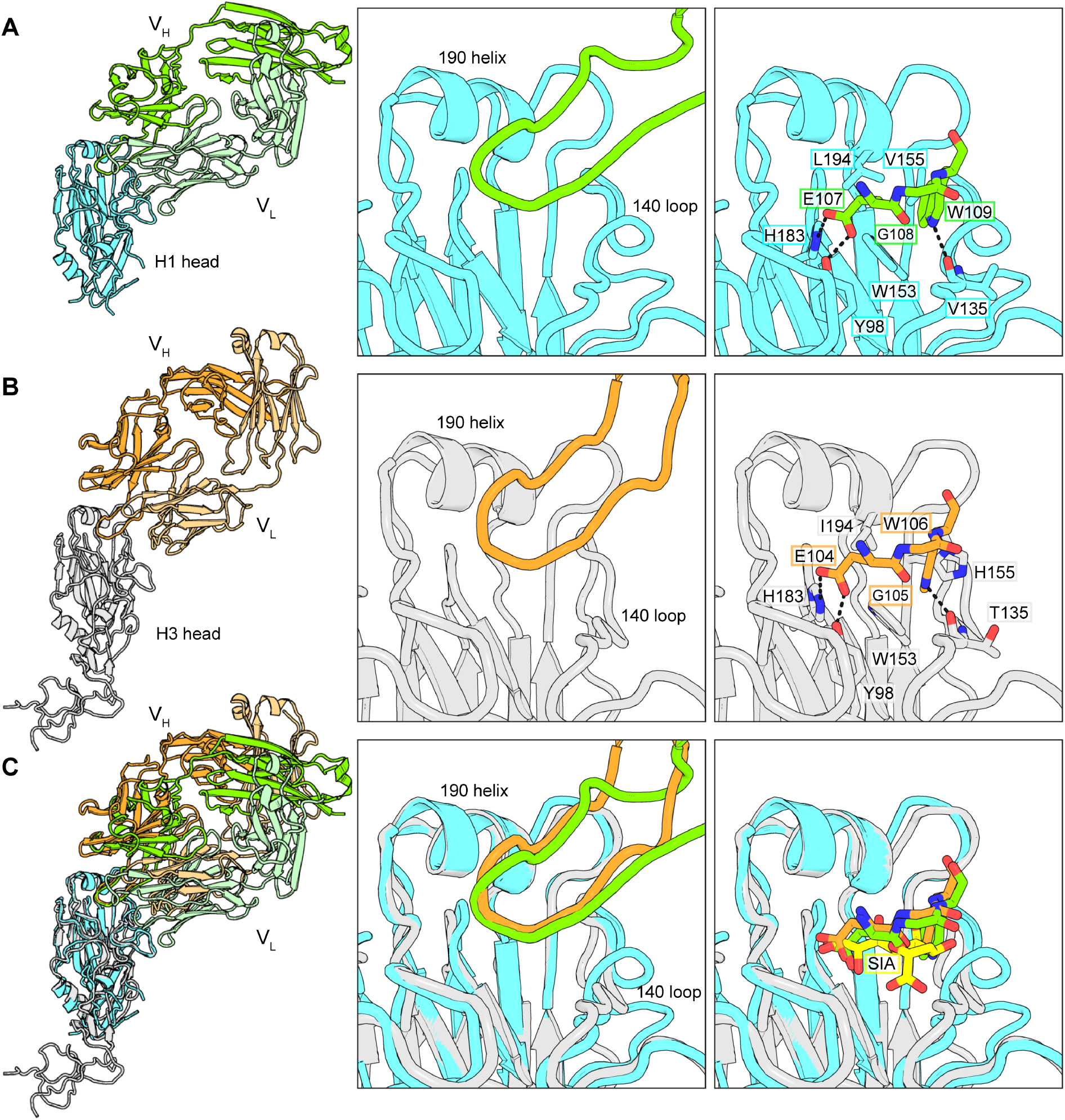
Structures of K03.28 and S8V1-172 Fabs bound to HA. **A**. Structure of K03.28 Fab bound with the HA head domain of A/California/07/2009 (H1N1)(NYMC-X181) (H1-X-181). The left panel shows the full structure, the middle panel the insertion of HCDR3 into the HA receptor binding site, and the right panel shows HCDR3 contacts that mimic those made by sialic acid. The K03.28 heavy chain is colored green, and its light chain is colored light green. The H1-X-181 HA head domain is colored cyan. Key features are indicated with labels. **B**. Structure of S8V1-172 Fab bound with the HA head domain of A/Sydney/05/1997(H3N2) (H3-SYD-1997) The right panel shows the full structure, the middle panel the insertion of HCDR3 into the HA receptor binding site and the right panel shows HCDR3 contacts that mimic those made by sialic acid. The S8V1-172 heavy chain is colored orange, and its light chain is colored light orange. The H3-SYD-1997 HA head domain is colored cyan. Key features are labeled. **C**. A superposition on the HA head domain of panels B and C. A sialic acid (yellow) in the receptor binding site has been added for comparisons using PDB 3UBE.

### A common sialic-acid mimicking motif

Donors S1, S5, S8, and S9 received the 2015-2016 trivalent inactivated seasonal influenza vaccine (Fluvirin), containing HAs from A/California/07/2009/H1N1(X-157), A/South Australia/55/2014(H3N2)(IVR-175), and B/Phuket/3073/2013 (B Phuket)^22^. HA specific Bmem cells from peripheral blood mononuclear cells (PBMCs) were obtained prior to immunization (visit one: V1) and seven days post immunization (V2).

We queried sequences from these circulating Bmem cells for additional antibodies with an HCDR3 Glu-Gly-Trp motif. We identified 10 examples from two donors S5 (n=2) and S8 (n=8) (Figure S2). The two antibodies from S5 are clonally related while the eight antibodies from S8 arose from at least four clonal B cell lineages (two light chain sequences were not recovered). Among these 10, the heavy chains derived from three VH genes, and the light chains and were paired with lambda or kappa light chains. Each has a 17 amino acid HCDR3, encoded in part from an IGHJ6 gene segment (Figure S2).

Antibody S8V1-172 typifies these IgGs. It is clonally related to at least two additional antibodies from S8 that share the Glu-Gly-Trp motif (S8V2-144 and S8V2-67), making it the largest lineage among the 10 antibodies with this motif. Except for the common IGHJ6, all antibody genes in that set are different from the K03.28 genes, including kappa rather than lambda light chains. K03.28 and S8V1-172 also have CDR3s of different lengths, 19 versus 17 for HCDR3 and 11 versus 8 for LCDR3, respectively (Figure S2).

We determined the structure of S8V1-172 complexed with the A/Sydney/05/1997(H3N2) HA head domain and compared it with K03.28 (Figure 2B). The pose of both antibodies is similar, but not identical (Figure 2C). With reference to the HA head, S8V1-172 is pitched toward the 190-helix, contacting it with its heavy chain. K03.28 is pitched away from the 190-helix, with minimal contacts. As a result, the HCDR3s of each antibody emanate from different points above the RBS but converge to closely related contacts for the Glu-Gly-Trp motif. As in K03.28, Glu-104 has polar contacts with Tyr-98 and His-183, Gly-105 has a positive phi angle, and Trp-106 mimics contacts by the sialic acid acetomido group. Receptor mimicry by this motif produces sufficient space to accommodate diversity at HA residue 226, a position that influences receptor specificity and antigenicity (Figure 2C). In H1s, 226 is typically Gln (or Arg in egg-propagated viruses), but it varies in circulating H3s (Ile, Leu, Gln, and Val).

**Figure 2.**
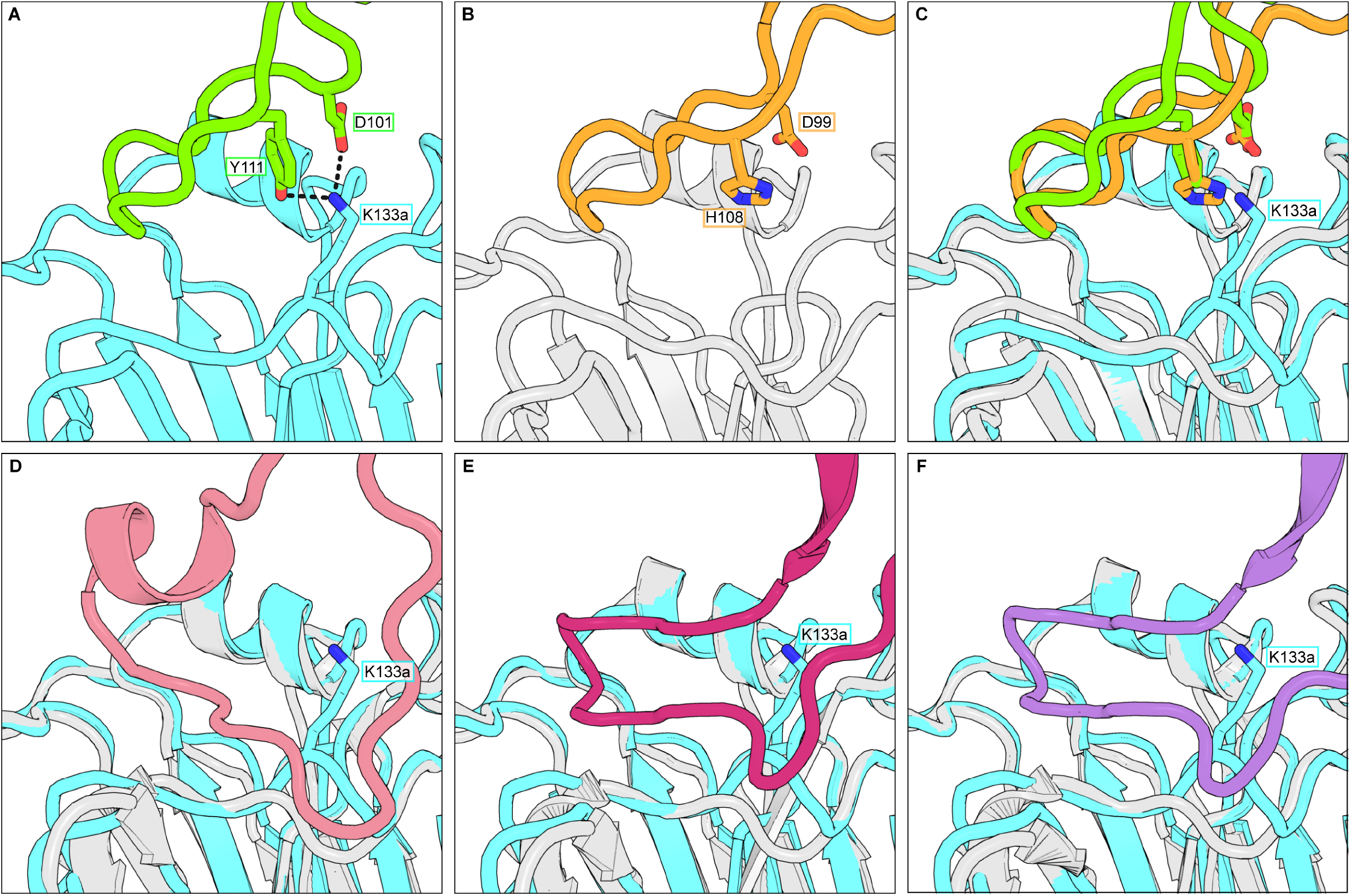
Accommodation of residue 133a. Antibodies K03.28 and S8V1-172 avoid the 133a bulge and engage Lys-133a. **A**. The HCDR3 from K03.28 (green) engaging the RBS of an H1-X-181 HA head domain (cyan). **B**. The HCDR3 from S8V1-172 (orange) engaging the RBS of an H3-Syd-97 HA head domain (gray). **C**. A superposition on the HA head of panels A and B. **(D-F)**. Superpositions between the H1-X-181 HA head domain (cyan) and H3 HA head domains (gray) bound with previously described H1-H3 cross neutralizing antibodies. These antibodies clash with the 133a bulge and the Lys sidechain. **D**. F045-92 (pink) (PDB 4O58) ^5^, **E**. C05 (magenta) (PDB 4FP8)^11^. **F**. K03.12 (purple) (PDB 5W08)^8^. Key residues shown in sticks are labeled throughout.

### Accommodation of K133a

The Fab-HA complexes of K03.28 and S8V1-172 have closely related structural adaptations to the K133a insertion. Their HCDR3s insert into the RBS at an angle that elevates them well above any potential clashes with 133a (Figure 2A-2C). In contrast, the HCDR3s of potent H1-H3 neutralizing antibodies C05^11^, K03.12^8^, and F045-092^5^, have extensive contacts along a strand spanning HA residues 130-138 and would clash with the 133a induced Cα bulge (Figure 2D-2E). The side chain of Lys at position 133a, present in seasonal H1s, would also introduce considerable interference. Rather than evading K133a, antibodies K03.28 and S8V1-172 engage it using analogous HCDR3 contacts (Figure 2A-2C). K03.28 coordinates K133a, present in the structure, with HCDR3 residues Asp-101 and Tyr-111 (Figure 2A). The structure of S8V1-172 with an H3 HA, which lacks 133a, suggests that it could engage K133a in an H1 HA with HCDR3 residues Asp-99 and His-108 (Figure 2C).

### Binding

We determined by enzyme-linked immunosorbent assays (ELISAs) the breadth of binding of K03.28 and S8V1-172 to a panel of historic H1N1 and H3N2 HAs (Figure 3). Both engaged HAs from each serotype. While K03.28 was broader, both bound H3 HAs from 1994 and 1997 and H1 HAs from before and after the 2009 pandemic. S8V1-172 bound only those H1 HAs with the Q226R mutation, which is quite common in vaccine strains (e.g., A/Solomon Islands/03/2006(H1N1) and A/California/07/2009(H1N1)(X-181) in the present study). For K03.28, binding to H1 HAs that circulated after 2015 also appears to be R226 dependent, as it bound A/Brisbane/02/2018 (H1N1)(IVR-190), A/Victoria/2570/2019 (H1N1)(IVR-215), both with R226, while failing to bind A/Wisconsin/588/2019(H1N1) with Q226.

**Figure 3.**
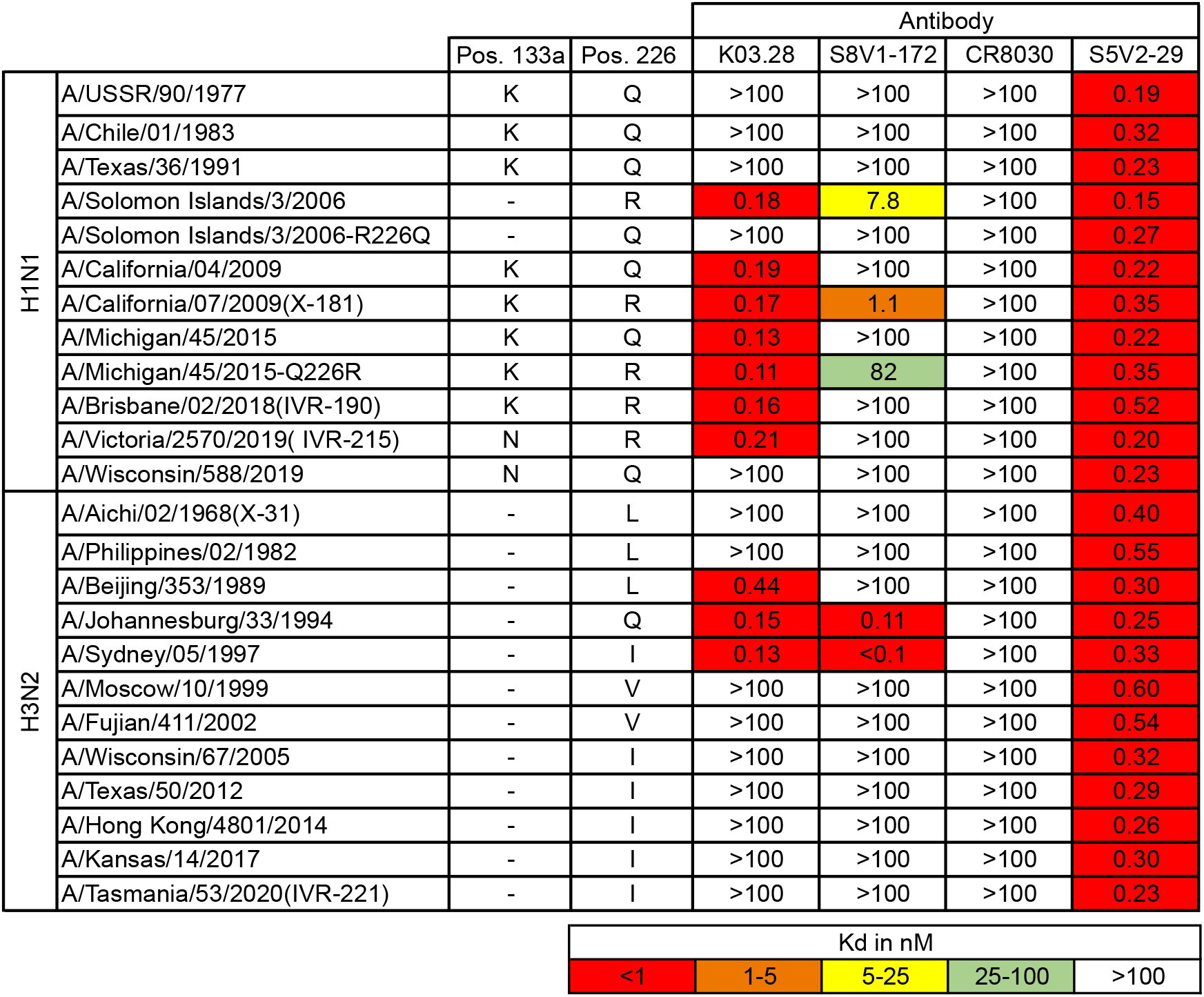
Breadth of binding by K03.28 and S8V1-172. Dissociation constants from ELISA measurements. The broad-influenza A virus-binding S5V2-29^22^ antibody was used as a positive control, and an influenza B virus-specific RBS-directed antibody, CR8033^25^, was used as a negative control for influenza A virus isolates. The presence and identity of residue a position 133a (Pos. 133a) and position 226 are indicated. Boxes are colored according to the key at the bottom. Mutations introduced into HAs are denoted.

We tested the R226 dependency by ELISA analysis of antibody binding to mutant HAs (Figure 3). We reverted the R226 in A/Solomon Islands/03/2006(H1N1) to Q226, which was present in circulating viruses. While K03.28 and S8V1-172 bound the R226 variant they failed to bind the R226Q revertant. We also introduced Q226R into A/Michigan/45/2015(H1N1), a mutation that is present in egg-passaged A/Michigan/45/2015(H1N1) stocks and egg-grown vaccines. This mutation conferred binding by K03.28 and S8V1-172. Of the two mutations that distinguish A/Victoria/2570/2019 (H1N1)(IVR-215) from A/Wisconsin/588/2019(H1N1), the difference at position 226 is the only one that falls within the epitopes of either antibody. Thus, the H1 Q226R mutation modulated binding by these antibodies in multiple distinct antigenic backgrounds.

### A common signature of H1-H3 cross-reactive antibodies

Single B cell culture antibodies from S1, S5, S8 and S9 were screened for binding to a panel of HAs using multiplex Luminex assays^22^. Many of these H1 and H3 HAs were from the same strains as those used in the ELISA with K03.28 and S8V1-172. The Luminex panel also included trimerized HA head domains from A/Johannesburg/33/1994(H3N2) and other H3 HAs. We searched those data for antibodies that shared a pattern of reactivity with K03.28 and S8V1-172.

Of 449 influenza A or B reactive Bmem cells, 39 (8.7%), had a pattern of antibody reactivity similar to that of K03.28 and S8V1-172. This group included all 10 antibodies with a HCDR3 Glu-Gly-Trp motif (Figure 4A). Like K03.28, two of the ten also bound the HA of circulating A/California/04/2009(H1N1), indicating that their engagement of H1 and H3 HAs did not depend on the Q226R mutation in this HA. Reactivity of the remaining 29 antibodies was virtually indistinguishable from that of the 10 with a Glu-Gly-Trp motif (Figure 4B), although they had not been identified in our initial motif search. Among these antibodies, we found no strong genetic biases, either in V(D)J or lambda and kappa loci, except for frequent IGHJ6 (90%) usage. We produced an alignment and HCDR3 sequence logo to identify commonalities (Figure S2). A central enrichment of Gly-Glu-Gly-Trp residues was prominent. Circulating Bmem with the Glu-Gly-Trp motif or related derivatives appear to be widespread and to associate with H1-H3 cross reactivity.

**Figure 4.**
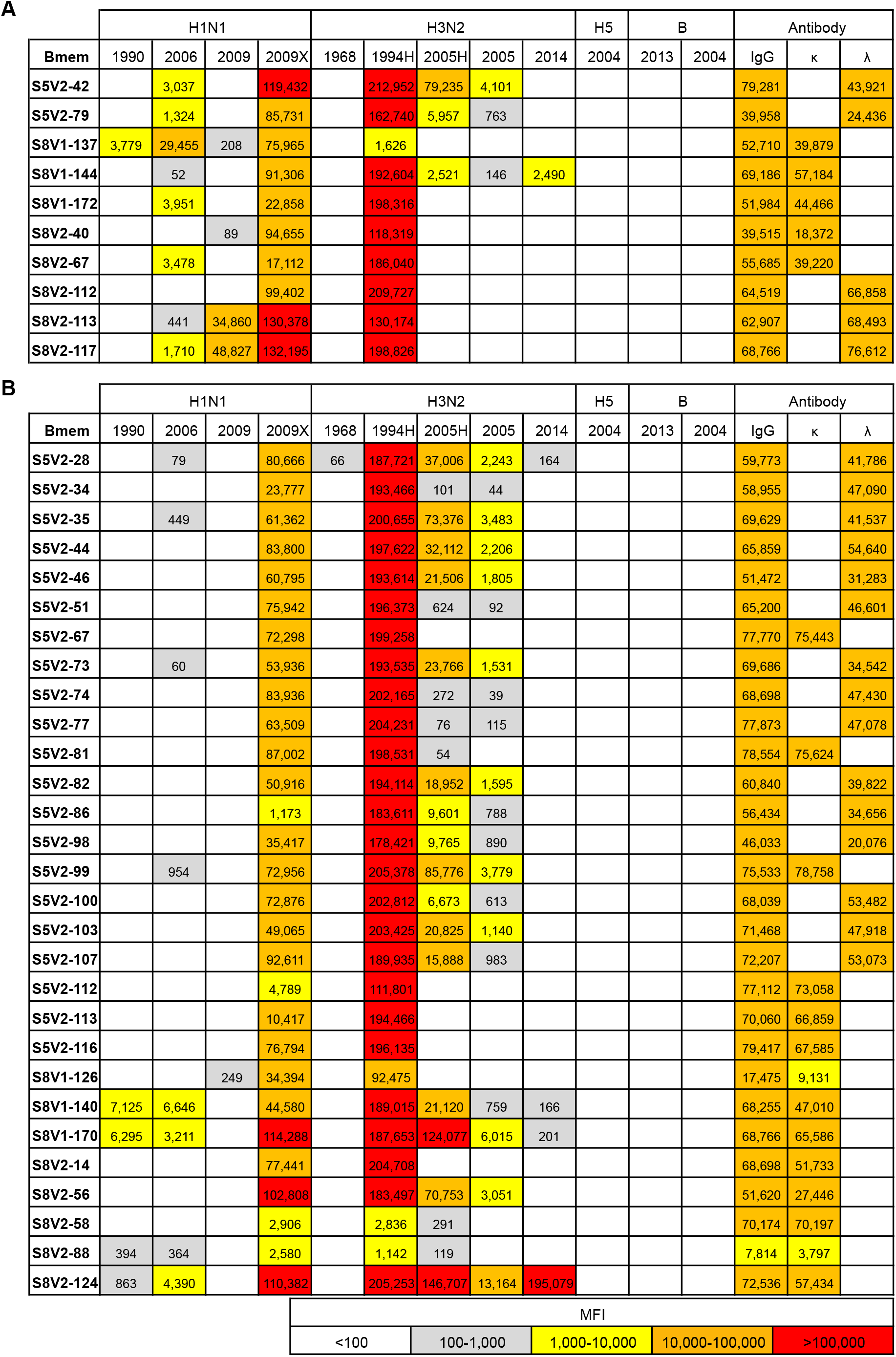
Common sequence motifs underlie common patterns of HA reactivity. Antibody reactivity for the 10 antibodies with a Glu-Gly-Trp motif **(A)** and the 29 antibodies that reacted similarly with the HAs in panel **(B)**. MFI values from multiplex bead assay for indicated antigens, colored according to the key. The HAs listed by year correspond to: H1N1 isolates: 1990 (A/Massachusetts/1/1990(H1N1)), 2006 (A/Solomon Islands/03/2006(H1N1), 2009 (A/California/04/2009(H1N1), 2009X (California/07/2009(H1N1)(X-181). H3N2 isolates: 1968 A/Aichi/2/1968(H3N2)(X-31-68), 1994H trimerized HA head domain from A/Johannesburg/33/1994(H3N2), 2005 A/Wisconsin/67/2005(H3N2), 2005H trimerized HA head domain from A/Wisconsin/67/2005(H3N2), 2014 A/South Australia/55/2014(H3N2). H5 isolate: 2004 H5N1 (A/Vietnam/1203/2004). B isolates: 2013 (B/Phuket/3073/2013) and 2004 (B/Malaysia/2506/2004). Antibodies specific for human IgG, lambda, and kappa chains were used for detection.

Of the 39 antibodies that bound H1 and H3 HAs, 34 reacted specifically with H1 HAs with the Q226R mutation (vaccine strains A/Solomon Islands/03/2006(H1N1) and A/California/07/2009(H1N1)(X-181)) and failed to bind H1 HAs with Q226 (circulating A/California/04/2009(H1N1) and A/Massachusetts/01/1990(H1N1)) (Figure 4). Most of these antibodies (31 of 34) were recovered following a documented immunization with an egg-propagated vaccine known to have the Q226R mutation. This set included all 23 antibodies from donor S5, born between 1990 and 1995^22^, with reactivity similar to K03.28 and S8V1-172. Most of the R226 specific antibodies from S8 were also recovered after receipt of the same vaccine. Only 5 of the 35 antibodies reacted with the H3 component of the vaccine (A/South Australia/55/2014(H3N2)(IVR-175)) and just one bound strongly. The remaining 30 instead bound most avidly to HAs of older isolates, often A/Johannesburg/33/1994(H3N2). These observations suggest that the egg-adapted H1 HA component of the vaccine interacted with Bmem cells that were elicited by prior exposure to an H3 virus.

We produced germline reverted antibodies for the S8V1-172 lineage and a 15-membered lineage from donor S5 (Figure S4). These inferred unmutated common ancestors (UCAs) represent the most probable BCR gene sequences of the naive B-cell progenitors. All non-germline nucleotides were identical in the S8V1-172 lineage but with two potential V-J recombination sites in the light chain and designated S8-UCA-1 and S8-UCA-2. The identities of non-germline nucleotides in the progenitor of the S5 lineage were inferred from a lineage phylogeny. S8-UCA-1 bound with detectable affinities both to A/Johannesburg/33/1994 (H3N2) and to the vaccine strain A/California/07/2009(H1N1)(X-181) with the Q226R mutation (Figure S5). No interaction was detected for circulating A/California/04/2009(H1N1) with Q226. The S5 UCA bound strongly with A/Johannesburg/33/1994(H3N2) and A/Sydney/05/1997(H3N2). For these lineages, updating of H3-specific immunity by vaccines containing R226 H1 HAs likely produced the H1 responses observed in the Bmem we sampled and characterized.

## Discussion

Engagement of cell-surface sialic acid moieties is an obligate step in the influenza virus replicative cycle^13^. Broadly neutralizing antibodies have been identified that mimic its contacts, while minimizing those with poorly conserved residues ^2,3,5-8,11,23,24 25,26^. The antibodies described here engage sialic acid coordinating residues with a novel sequence motif that is abundant in the human B memory cell compartment. Antibodies belonging to this class cross-neutralize H1 and H3 viruses, agnostic to the presence or absence of the 133a insertion. Surmounting this steric barrier distinguishes these antibodies from previously described H1-H3 neutralizing antibodies^5,8,11^.

Members of this antibody class have a common pattern of HA reactivity, defined by avid engagement of H3 HA isolates from the 1990s and H1 HAs from the 2009 H1N1 pandemic. Over a decade separates circulation of these viruses, and most of the antibodies do not engage HAs from intervening years. The pattern suggests that sequential H3 and H1 exposures gave rise to the observed lineages, especially in view of the differences among the donors and the absence of any common IGHV or IGVK/L gene usage. The response was particularly robust in S5 and S8, both of whom were infants or small children in the early 1990s. Infection with an early 1990s strain was therefore probably their first influenza exposure and hence the source of their initial immune imprint, as pediatric vaccination was not yet recommended at that time. KEL03, born in 1975, could plausibly have experienced an infection with an H3N2 virus during the 1990s and probably received an H3N2 vaccination. All three donors were likely susceptible to infection 2009 H1N1 pandemic viruses, and each did indeed receive H1N1 vaccinations after 2009.

When confident inference of the UCA of a lineage is possible, then the reactivity of that UCA with a panel of historical strains can identify the likely date of the primary exposure that gave rise to that lineage^24^. The uniquely determined UCA of the lineage from S5 and one of the two possible UCAs of the lineage from S8 both bind HA from A/Johannesburg/33/1994 (H3N2), but not much earlier or much later H3 HAs (Table in Fig. S5), as expected for a primary exposure in the early 1990s. Although K03.28 has no clonal lineage siblings in our data set, its reactivity also suggests that it derives from an early 1990s primary response. Since the K03 donor was born almost 20 years earlier, the properties of the antibody suggest that even if recall of memory from a first exposure appears to dominate later responses (i.e. “immune imprinting” ^27,28^), primary responses to later exposures can contribute new specificities to the Bmem repertoire.

Most administered influenza vaccines deliver HA immunogens derived from viruses propagated in embryonated eggs. Egg growth can yield RBS mutant viruses with tropism for cells bearing α2,3 sialic acids (required for avian transmission) instead of or in addition to α2,6-linked sialic acids (required for human transmission)^13,14^. The Q226R mutation is a characteristic H1 substitution and frequently occurs in new pandemic vaccines ^15,18,29^ While not absolute, many antibodies described here depend upon R226 for H1 binding. Immunization with H1 R226 HAs has been shown to misdirect H1 antibody responses^15,18^. Our observations suggest that R226 can also influence H3 and H1-H3 responses. Whether there is some relationship between the RBS antigenic surfaces of mid-1990s H3 HAs and the initial 2009 pandemic vaccine A/California/07/2009(H1N1)(X-181) HA, as the observations from S5 and S8 suggest, will need additional examples. Nonetheless, in view of the observation that a single residue can govern H1-H3 neutralization, barriers to cross-serotype neutralization by RBS-directed antibodies may be relatively low. These and previous results also illustrate the need to transition away from egg-grown components (or even egg-selected reassortants) in influenza vaccine production.

Conserved sialic acid-coordinating HA residues provide a target for broadly neutralizing antibodies^2,3,5-8,11,23,24 25,26^. Broadly protective antibodies directed to the HA-stem or HA-head interface generally require antibody Fc-mediated killing of already infected cells^22,30,31^, while genuinely neutralizing antibodies block the apparent establishment of infection. Although individual RBS directed antibodies lack universal breadth, their abundance and lack of genetic restrictions suggest that polyclonality can achieve broad protection. Antibodies K03.28 and those from the K03.12 lineage were isolated from donor KEL03^8^. Together they engage H1 HAs from 1995-2015 and nearly all H3 HAs from 1994 to at least 2014. This breadth includes pre- and post-2009 pandemic H1N1 isolates. Achieving broadly protective immunity will likely require polyclonal responses, directed to multiple conserved epitopes and that protect by multiple mechanisms. The antibodies described here expand the potential repertoire of broadly neutralizing antibodies that can contribute to such broad protection.

## Methods

### Cell lines

Human 293F cells were maintained at 37°C with 5% CO2 in FreeStyle 293 Expression Medium (ThermoFisher) supplemented with penicillin and streptomycin. High Five™ Cells (BTI-TN-5B1-4) (Trichoplusia ni) were maintained at 28°C in EX-CELL 405 medium (Sigma) supplemented with penicillin and streptomycin.

### Recombinant Fab expression and purification

Synthetic heavy- and light-chain variable domain genes for Fabs were cloned into a modified pVRC8400 expression vector, as previously described^32^. Fab fragments used in crystallization were produced with a C-terminal, noncleavable 6xhistidine (6xHis) tag. Fab fragments were produced by polyethylenimine (PEI) facilitated, transient transfection of 293F cells that were maintained in FreeStyle 293 Expression Medium. Transfection complexes were prepared in Opti-MEM and added to cells. Supernatants were harvested 4-5 days post transfection and clarified by low-speed centrifugation. Fabs were purified by passage over Co-NTA agarose (Clontech) followed by gel filtration chromatography on Superdex 200 (GE Healthcare) in 10 mM Tris-HCl, 150 mM NaCl at pH 7.5 (buffer A).

### Recombinant IgG expression and purification

The heavy chain variable domains of selected antibodies were cloned into a modified pVRC8400 expression vector to produce a full length human IgG1 heavy chain^22,32,33^. IgGs were produced by transient transfection of 293F cells as specified above. Five days post-transfection supernatants were harvested, clarified by low-speed centrifugation, and incubated overnight with Protein A Agarose Resin (GoldBio). The resin was collected in a chromatography column, washed with a column volume of buffer A, and eluted in 0.1M Glycine (pH 2.5) which was immediately neutralized by 1M tris(hydroxymethyl)aminomethane (pH 8.5). Antibodies were then dialyzed against phosphate buffered saline (PBS) pH 7.4.

### Recombinant HA expression and purification

Recombinant (HA) head domain constructs were expressed by infection of insect cells with recombinant baculovirus as previously described^32,33^. In brief, a synthetic DNA corresponding to the full-length the globular HA-head was subcloned into a pFastBac vector modified to encode a C-terminal rhinovirus 3C protease site and a 6xHis tag. Supernatant from recombinant baculovirus infected High Five™ Cells (Trichoplusia ni) was harvested 72 hr post infection and clarified by centrifugation. Proteins were purified by adsorption to Co-NTA agarose resin, followed by a wash in buffer A, a second wash (trimers only) with buffer A plus 5-7mM imidazole, elution in buffer A plus 350mM imidazole (pH 8) and gel filtration chromatography on a Superdex 200 column (GE Healthcare) in buffer A.

Full-length HA ectodomain (FLsE) were produced by polyethylenimine (PEI) facilitated, transient transfection of 293F cells maintained in FreeStyle 293 Expression Medium. Synthetic DNA corresponding to the full-length ectodomain (FLsE) were cloned into a pVRC vector modified to encode a C-terminal thrombin cleavage site, a T4 fibritin (foldon) trimerization tag, and a 6xHis tag^32,34^. Transfection complexes were prepared in Opti-MEM and added to cells. Supernatants were harvested 4-5 days post transfection and clarified by low-speed centrifugation. HA trimers were purified by passage over Co-NTA agarose (Clontech) followed by gel filtration chromatography on Superdex 200 (GE Healthcare) in 10 mM Tris-HCl, 150 mM NaCl at pH 7.5 (buffer A).

### ELISA

Five hundred nanograms of rHA FLsE were adhered to high-capacity binding, 96 well-plates (Corning) overnight in PBS pH 7.4 at 4°C. Plates were washed with a PBS-Tween-20 (0.05%v/v) buffer (PBS-T) and then blocked with PBS-T containing 2% bovine serum albumin (BSA) for 1 hr at room temperature. Blocking solution was then removed, and 5-fold dilutions of IgGs (in blocking solution) were added to wells. Plates were then incubated for 1 hr at room temperature followed by removal of IgG solution and three washes with PBS-T. Secondary, anti-human IgG-HRP (Abcam ab97225) diluted 1:10,000 in blocking solution, was added to wells and incubated for 30 min at room temperature. Plates were then washed three times with PBS-T. Plates were developed using 150μl 1-Step ABTS substrate (ThermoFisher, Prod#37615). Following a brief incubation at room temperature, HRP reactions were stopped by the addition of 100μl of 1% sodium dodecyl sulfate (SDS) solution. Plates were read on a Molecular Devices SpectraMax 340PC384 Microplate Reader at 405 nm.

KD values for ELISA were obtained as follows. All measurements were performed in technical triplicate. The average background signal (no primary antibody) was subtracted from all absorbance values. Values from multiple plates were normalized to the S5V2-29 standard that was present on each ELISA plate. The average of the three measurements were then graphed using GraphPad Prism (v9.0). KD values were determined by applying a nonlinear fit (One site binding, hyperbola) to these data points.

### Virus microneutralization assays

Virus neutralization endpoint titers were determined using the influenza microneutralization assay as described^8,35-38^. Serum samples were heat inactivated for 60 minutes at 56°C and diluted 1:10 in virus diluent. Each sample was then diluted twofold in virus diluent yielding a range of 1:10 - 1:1280 in a flat-bottomed 96-well tissue culture plate. Samples were then mixed equimolar with virus diluent containing 100 TCID50 of each influenza virus of interest. Control samples included known antisera and naïve sera treated in exactly the same manner as experimental samples. After virus addition, samples are incubated for 60 minutes at 37°C 5% CO2. 1.5e4 MDCK cells (London strain, IRR FR-58) were added to each well. Plates were incubated overnight at 37°C 5% CO2. Each well was aspirated, and cells were washed one time with PBS. The PBS was aspirated. Then 250 µL of -20°C 80% acetone was added to each well, and plates were incubated at room temperature for 10 minutes. The acetone was removed and plates air-dried. Each well was washed three times with wash buffer. Primary antibody (Mouse Anti-Influenza A NP, Millipore MAB8251 or Mouse Anti-Influenza B NP, Millipore MAB8661) was diluted 1:4000 in antibody diluent, and 50 µL was added to every well. Plates were incubated for 60 minutes at room temperature. Each well was washed three times with wash buffer. Secondary antibody (Goat anti-mouse + horseradish peroxidase, KPL 474-1802) was diluted 1:4000 in antibody diluent, and 50 µL was added to every well. Plates were incubated for 60 minutes at room temperature. Each well was washed five times with wash buffer. 100 µL substrate was added to every well and incubated at room temperature. The reaction was stopped with 100 µL 0.5 N sulfuric acid after apparent color change was observed in virus-only control wells. Absorbance was read at 490 nM in a Synergy H1 automated microplate reader (BioTek Instruments Inc.). Wells with absorbance values less than or equal to 50% of virus-only control wells were scored as neutralization positive. Data were expressed as the geometric mean of the reciprocal of the final dilution factor that was positive for neutralization. All samples were assayed in at least duplicates.

Influenza viruses were propagated in embryonated specific-pathogen-free chicken hen eggs or MDCK(CCL-34) cells as described^8^. Reagents obtained through BEI Resources, NIAID, NIH include: Influenza A viruses A/Aichi/2/1968 (H3N2) NR-3177; Kilbourne F123: A/Victoria/3/1975 (HA, NA) x A/Puerto Rico/8/1934 (H3N2), Reassortant X-47 NR-3663; A/Philippines/2/1982 (H3N2) NR-28649; Kilbourne F178: A/Shanghai/11/1987 (HA, NA) x A/Puerto Rico/8/1934 (H3N2), High Yield, Reassortant X-99a NR-3505; Kilbourne F86: A/Johannesburg/33/1994 (HA, NA) x A/Puerto Rico/8/1934 (H3N2), Reassortant X-123a NR-3580; Kilbourne F97: A/Moscow/10/1999 (HA, NA) x A/Puerto Rico/8/1934 (H3N2), Reassortant X-137 NR-3587; polyclonal influenza virus, A/Aichi/2/1968 (H3N2) serum (guinea pig), NR-3126; NR-4282. A/USSR/90/1977, reassortant X-67 (H1N1) NR-3666; A/Chile/01/1983 reassortant X-83 (H1N1) NR-3585, A/Beijing/262/1995 reassortant X-127 (H3N2) NR-3571; A/Solomon Islands (H1N1) NR-41798.

Influenza A virus A/Wisconsin/67/2005 (H3N2), FR-397; and MDCK London cells (FR-58) were obtained through the International Reagent Resource (formerly the Influenza Reagent Resource), Influenza Division, WHO Collaborating Center for Surveillance, Epidemiology and Control of Influenza, Centers for Disease Control and Prevention, Atlanta, GA, USA.

### Crystallization

Fab fragments were co-concentrated with HA-head domains) at a molar ratio of ∼1:1.3 (Fab to HA-head) to a final concentration of ∼20 mg/ml. Crystals of Fab-head complexes were grown in hanging drops over reservoir solutions Crystals of the K03.28-A/California/07/2009(H1N1)(X-181) HA head domain were grown in hanging drops over a reservoir of 25% (w/v) poly(ethylene glycol) 1500. Crystals of S8V1-172 complexed with the HA head domain of A/Sydney/05/1997(H3N2) were grown in drops over a reservoir of 0.1 M tris(hydroxymethyl)aminomethane pH 8.5 and 25% (w/v) poly(ethylene glycol) 3350. Crystals were cryprotected with glycerol at concentrations of 22% in cryoprotectant buffers that were 20% more concentrated than the well solution. Cryprotectant was added directly to the drop, crystals were harvested, and flash cooled in liquid nitrogen.

### Structure determination and refinement

We recorded diffraction data at the Advanced Photon Source on beamline 24-ID-C. Data were processed and scaled (XSCALE) with XDS^39^. Molecular replacement was carried out with PHASER^40^, dividing each complex into four search models (HA-head, Vh, Vl and constant domain). Search models were 3UBE, 6E56, 4HK0, 4WUK for the K03.28-HA head domain complex and 6XPZ, 6E4X, 6MHR, 6E4X, for the S8V1-172-HA head domain complex. We carried out refinement calculations with PHENIX^41^ and model modifications, with COOT^42^. Refinement of atomic positions and B factors was followed by translation-liberation-screw (TLS) parameterization and, if applicable, placement of water molecules. All placed residues were supported by electron density maps and subsequent rounds of refinement. Final coordinates were validated with the MolProbity server^43^. Data collection and refinement statistics are in Table S1. Figures were made with PyMOL (Schrödinger, New York, NY).

## DATA AND SOFTWARE AVAILABILITY

Coordinates and diffraction data have been deposited at the PDB, accession numbers 7TRH and 7TRI.

## ACKNOWLEDGMENTS

We thank the many members of our Program Project Consortium for advice and discussion. We thank Lindsey R. Robinson-McCarthy for invaluable discussion and comments. X-ray diffraction data were recorded at beamline ID-24-C (operated by the Northeast Collaborative Access team: NE-CAT) at the Advanced Photon Source (APS, Argonne National Laboratory). We thank NE-CAT staff members for advice and assistance in data collection. NE-CAT is funded by NIH grant P30 GM124165. APS is operated for the DOE Office of Science by Argonne National Laboratory under contract DE-AC02-06CH11357. The research was supported by NIAID Program Project Grant P01 AI089618 (to G.H.K.) and funds from the University of Pittsburgh Center for Vaccine Research (to K.R.M).

## Supporting information

**Table S1.**
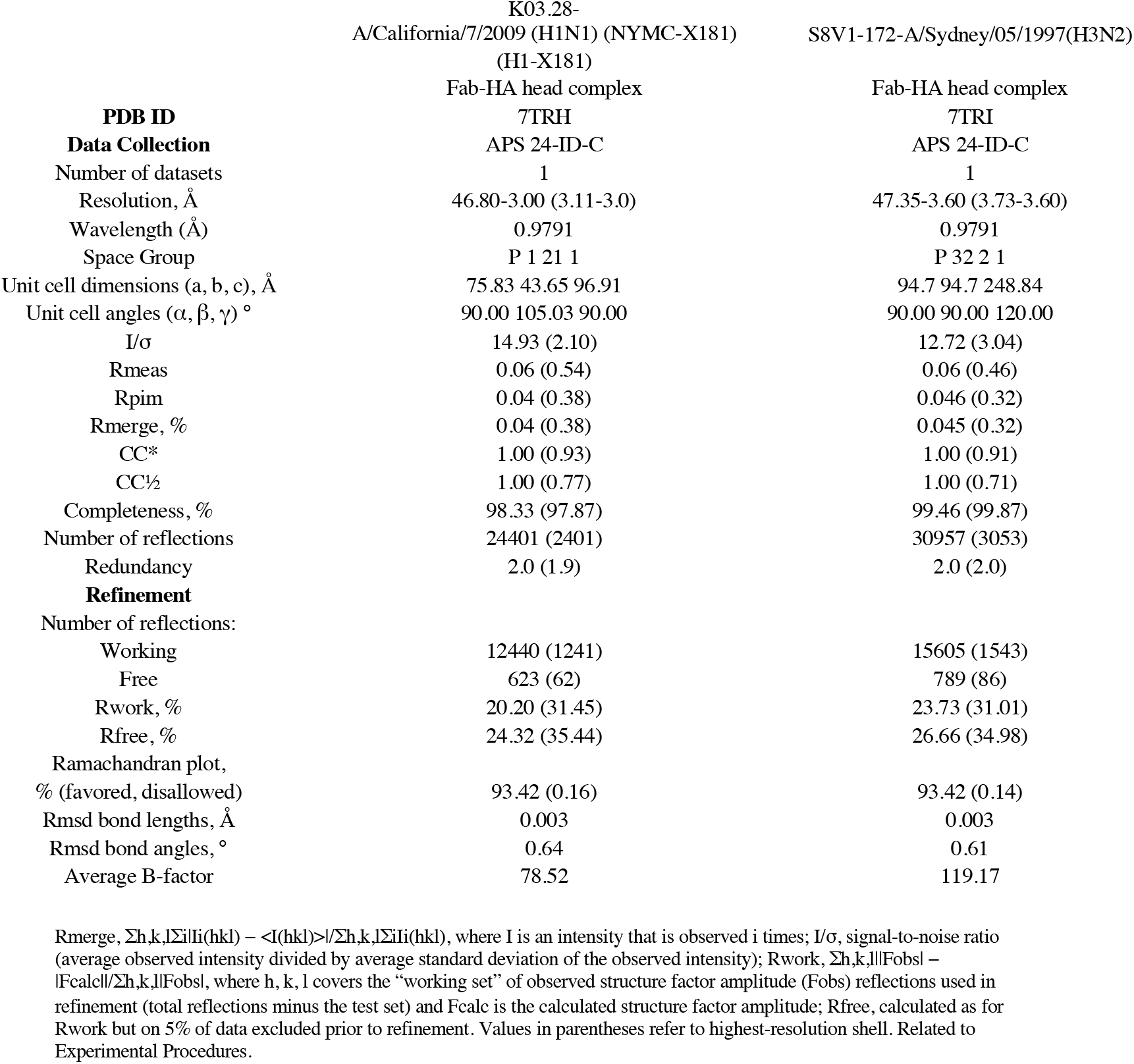
Data collection and refinement statistics.

**S1.**
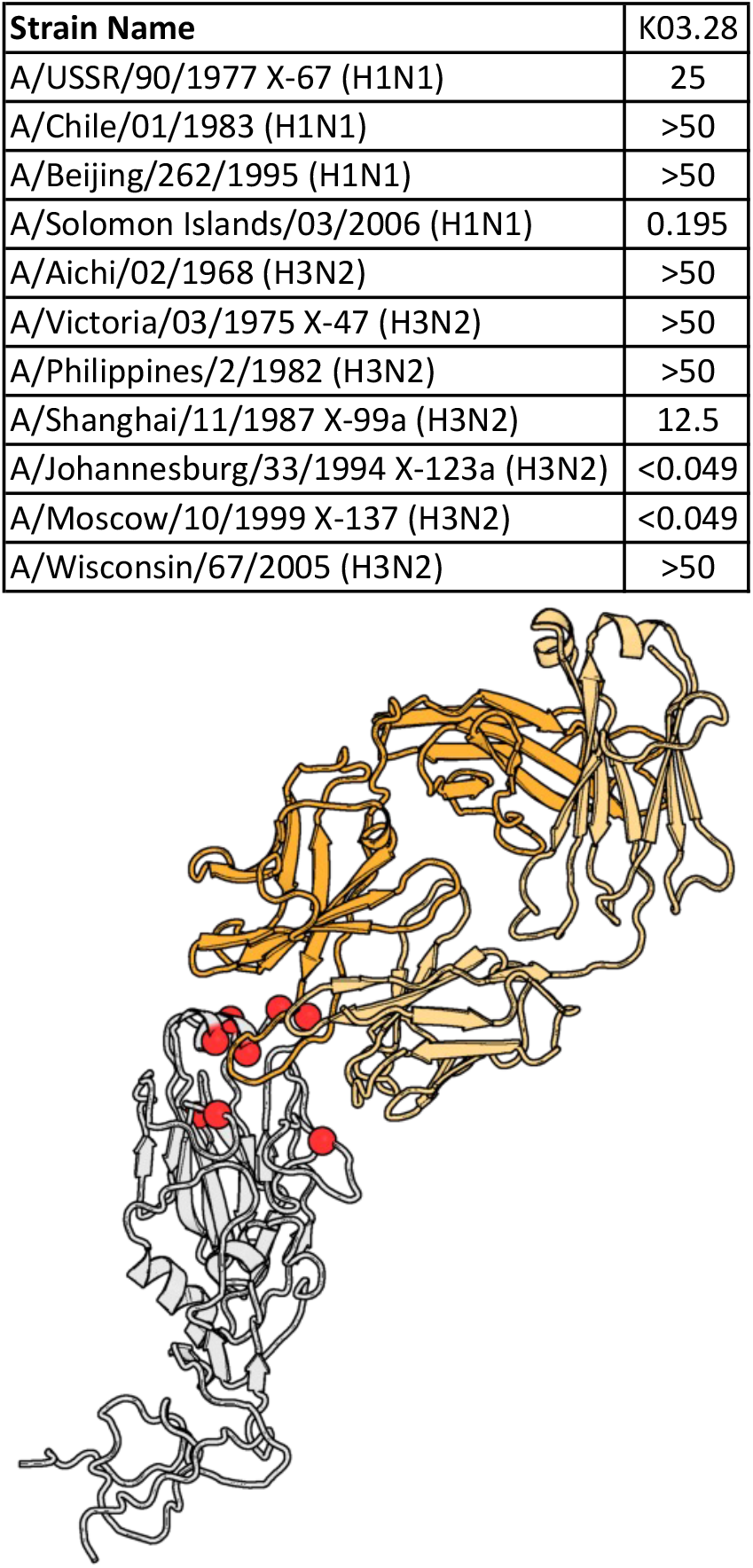
K03.28 microneutralization data. Top. Microneutralization titers for antibody K03.28. All values are in μg/ml. Bottom. Differences between the A/Moscow/10/1999(H3N2) (GenBank DQ487341) isolates used for HA binding assays and the A/Moscow/10/1999(H3N2)(X-137) reassortant virus used in microneutralization assays (GenBank CY121381). Eight amino acid substitutions are in the HA head domain and are predicted to impact antibody binding. Their positions are shown in red spheres in the structure of S8V1-172 complexed with the A/Sydney/05/1997(H3N2) HA head domain (145K/N, 158E/K,159N/Y, 190V/D, 194V/L 196V/T, 226I/V, and 246N/K.

**S2.**
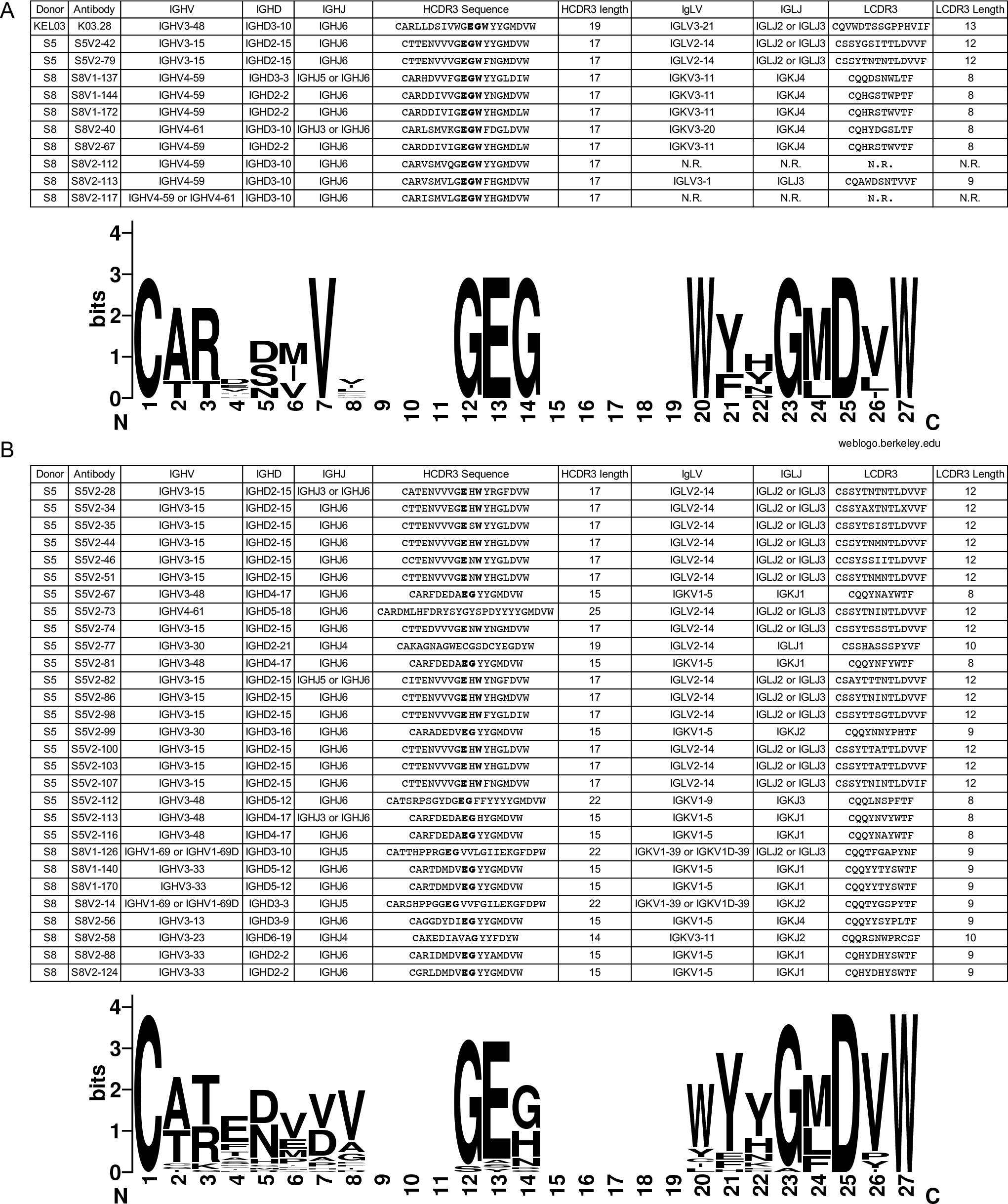
Antibody V(D)J usage and CDR3 sequences. Gene utilization and CDR3 sequences of the 10 antibodies with a Glu-Gly-Trp motif **(A)** or those with similar patterns of HA reactivity **(B)**. Amino acids in common with the Glu-Gly-Trp motif are bolded. Sequence logos below each panel were produced from an initial alignment of all antibody HCDR3 that was then subdivided into two alignments, one specific for each panel, without realignment. This was done to allow for comparisons between panel A and B. Gaps in the sequence logo are sites of length variation. Sequence logos were produced with WebLogo^44^.

**S3.**
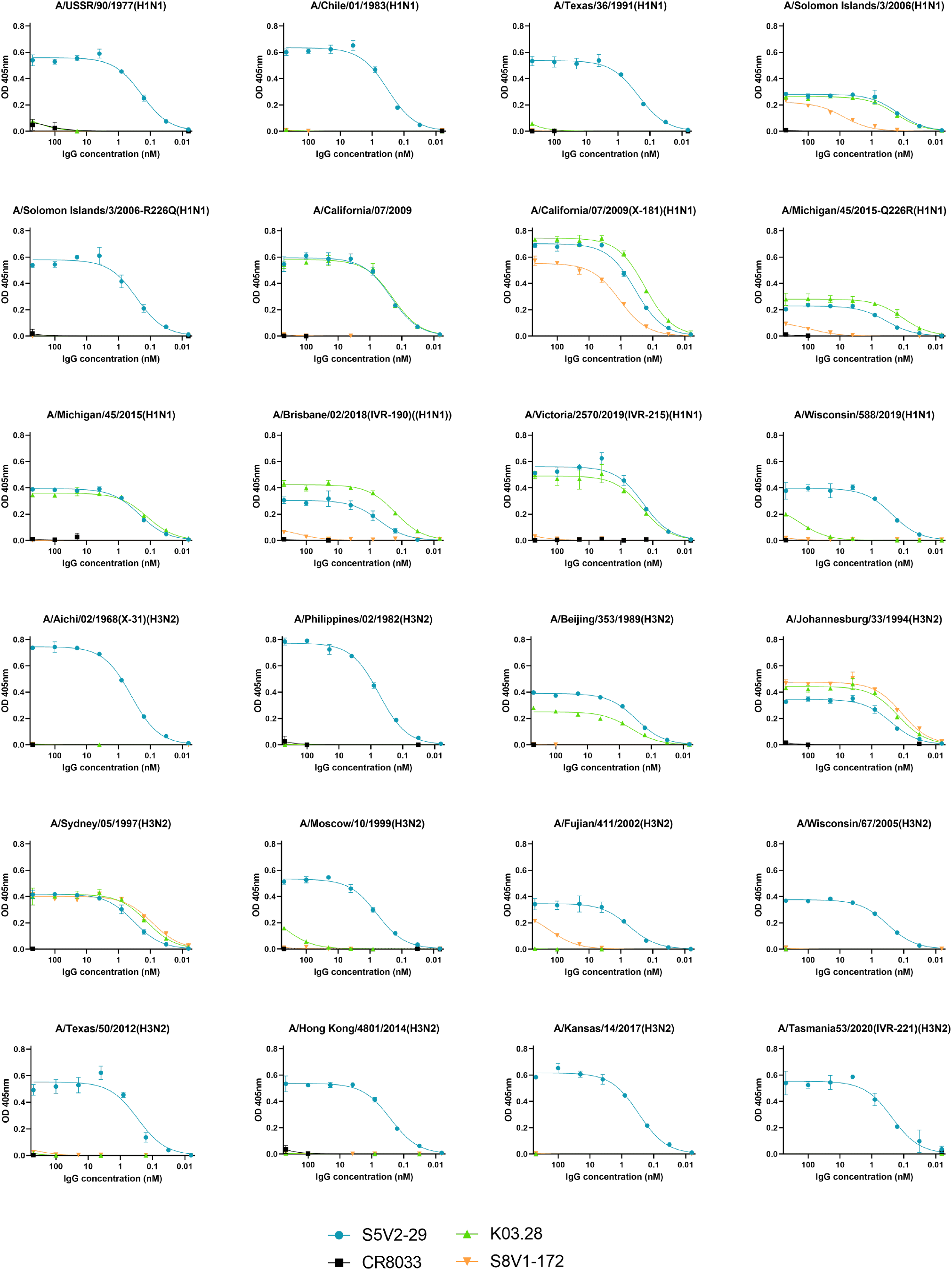
ELISA titrations of antibodies on HA coated plates. The broadly binding, head interface directed S5V2-29^22^ was used as a positive control and an influenza B specific, RBS directed antibody, CR8033^25^, as a negative control for influenza A isolates. Data points represent the average of three technical replicates. The standard error of the mean is shown for each point. KDs were calculated from the curves fit to these data points.

**S4.**
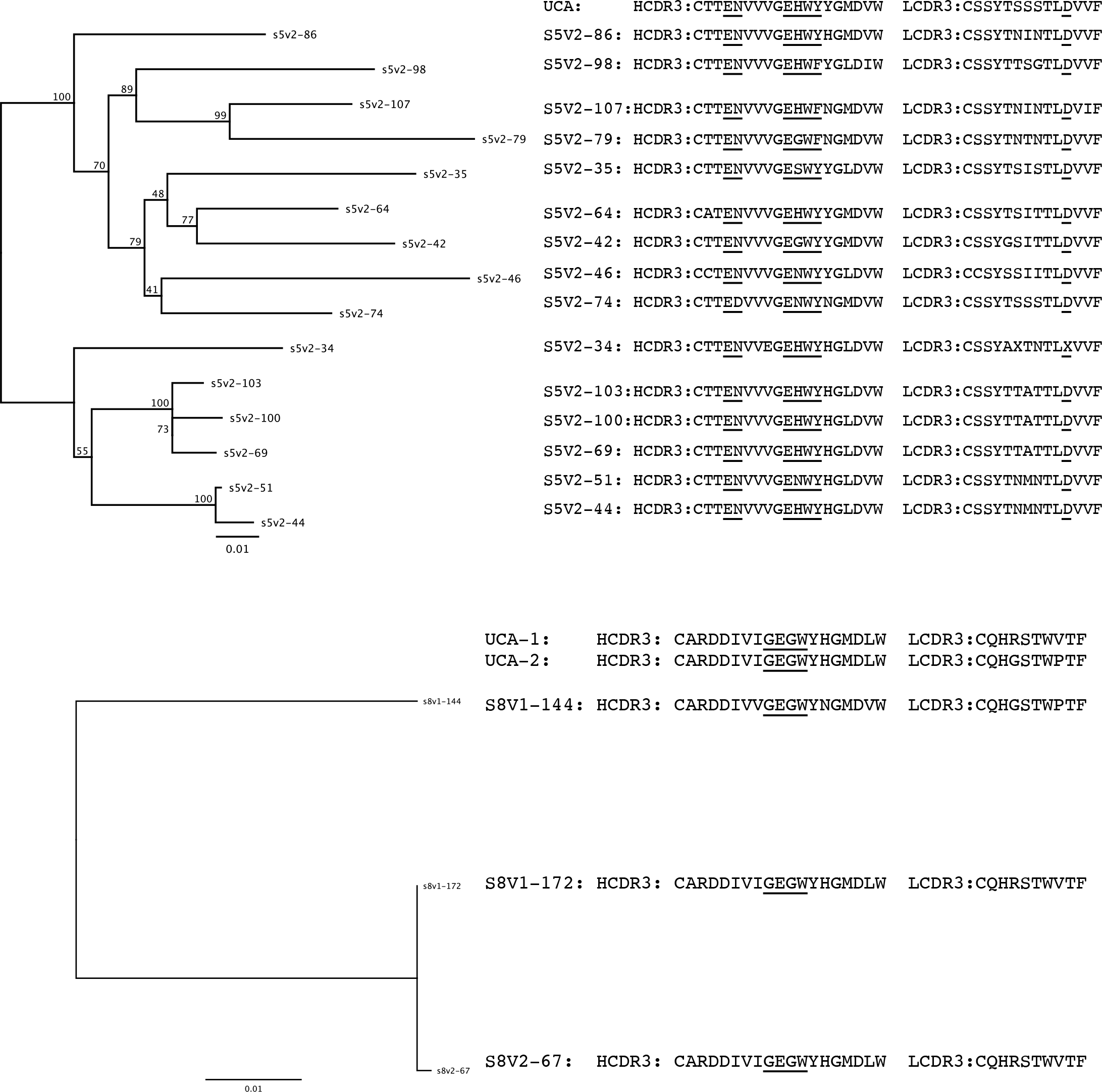
Germline reversion of select antibodies from clonal lineages. The phylogeny of two clonal antibody lineages are accompanied by amino acids sequences of each antibody’s CDR3. Amino acids encoded by N nucleotides are underlined.

**S5.**
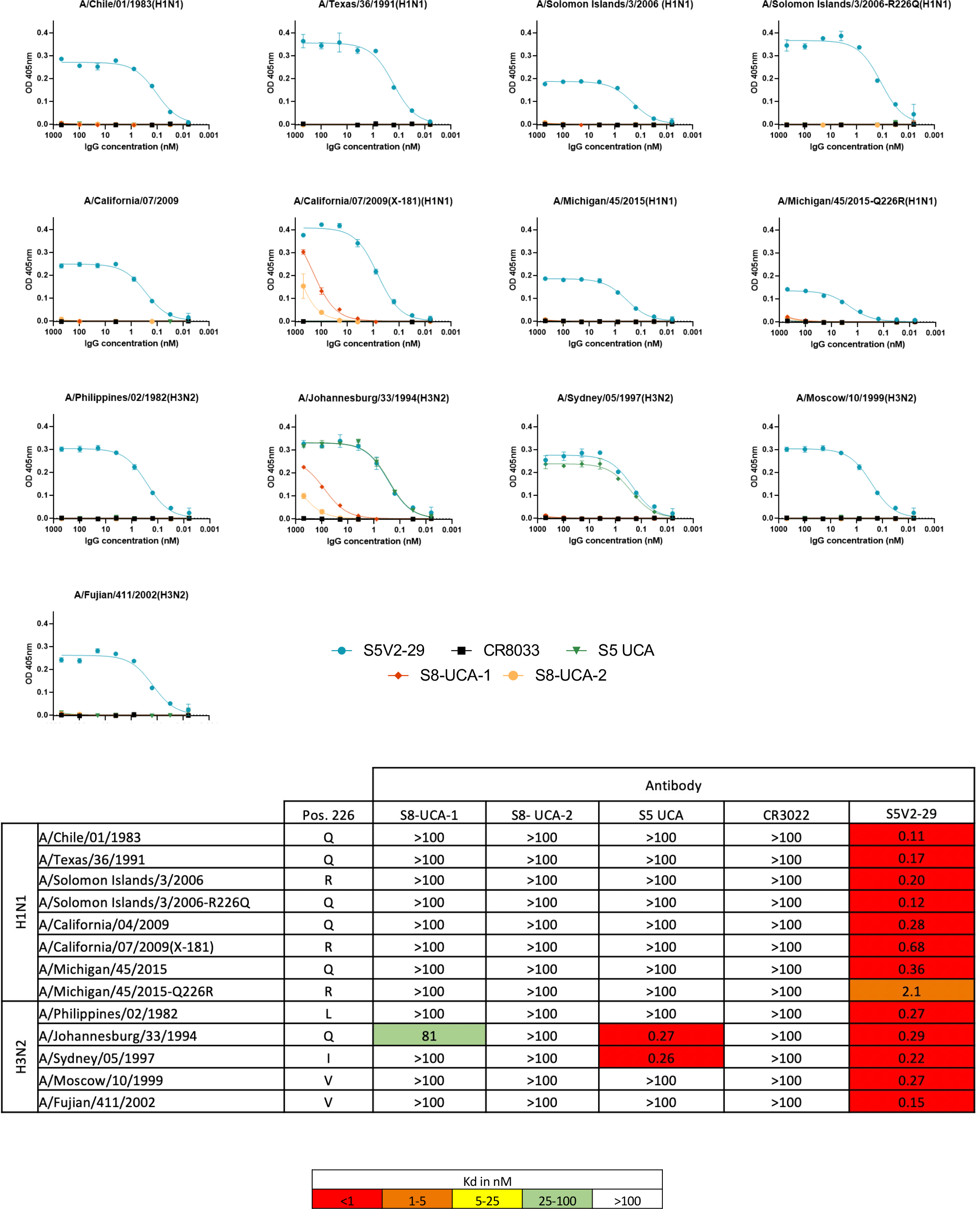
ELISA titrations and affinities of antibody UCAs. The broadly binding, head interface directed S5V2-29^22^ was used as a positive control and an influenza B specific, RBS directed antibody, CR8033^25^, as a negative control for influenza A isolates. Data points represent the average of three technical replicates. The standard error of the mean is shown for each point. KDs were calculated from the curves fit to these data points and are included below.

