## supporting figures for "A class of antibodies that overcome a steric barrier to cross-group neutralization of influenza viruses"

### Supporting Figure 1

| **Strain Name** | K03.28 |
|---|---|
| A/USSR/90/1977 X-67 (H1N1) | 25 |
| A/Chile/01/1983 (H1N1) | >50 |
| A/Beijing/262/1995 (H1N1) | >50 |
| A/Solomon Islands/03/2006 (H1N1) | 0.195 |
| A/Aichi/02/1968 (H3N2) | >50 |
| A/Victoria/03/1975 X-47 (H3N2) | >50 |
| A/Philippines/2/1982 (H3N2) | >50 |
| A/Shanghai/11/1987 X-99a (H3N2) | 12.5 |
| A/Johannesburg/33/1994 X-123a (H3N2) | <0.049 |
| A/Moscow/10/1999 X-137 (H3N2) | <0.049 |
| A/Wisconsin/67/2005 (H3N2) | >50 |

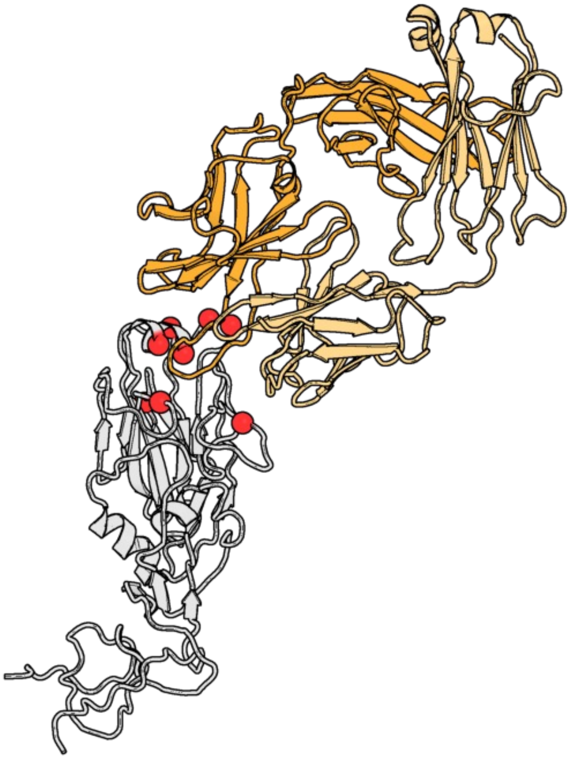

### Supporting Figure 2

A

| Donor | Antibody | IGHV | IGHD | IGHJ | HCDR3 Sequence | HCDR3 length | IgLV | IGLJ | LCDR3 | LCDR3 Length |
|---|---|---|---|---|---|---|---|---|---|---|
| KEL03 | K03.28 | IGHV3-48 | IGHD3-10 | IGHJ6 | CARLLDSIVWG**EGW**YYGMDVW | 19 | IGLV3-21 | IGLJ2 or IGLJ3 | CQVWDTSSGPPHVIF | 13 |
| S5 | S5V2-42 | IGHV3-15 | IGHD2-15 | IGHJ6 | CTTENVVVG**EGW**YYGMDVW | 17 | IGLV2-14 | IGLJ2 or IGLJ3 | CSSYGSITTLDVVF | 12 |
| S5 | S5V2-79 | IGHV3-15 | IGHD2-15 | IGHJ6 | CTTENVVVG**EGW**FNGMDVW | 17 | IGLV2-14 | IGLJ2 or IGLJ3 | CSSYTNTNTLDVVF | 12 |
| S8 | S8V1-137 | IGHV4-59 | IGHD3-3 | IGHJ5 or IGHJ6 | CARHDVVFG**EGW**YYGLDIW | 17 | IGKV3-11 | IGKJ4 | CQQDSNWLTF | 8 |
| S8 | S8V1-144 | IGHV4-59 | IGHD2-2 | IGHJ6 | CARDDIVVG**EGW**YNGMDVW | 17 | IGKV3-11 | IGKJ4 | CQHGSTWPTF | 8 |
| S8 | S8V1-172 | IGHV4-59 | IGHD2-2 | IGHJ6 | CARDDIVIG**EGW**YHGMDLW | 17 | IGKV3-11 | IGKJ4 | CQHRSTWVTF | 8 |
| S8 | S8V2-40 | IGHV4-61 | IGHD3-10 | IGHJ3 or IGHJ6 | CARLSMVKG**EGW**FDGLDVW | 17 | IGKV3-20 | IGKJ4 | CQHYDGSLTF | 8 |
| S8 | S8V2-67 | IGHV4-59 | IGHD2-2 | IGHJ6 | CARDDIVIG**EGW**YHGMDLW | 17 | IGKV3-11 | IGKJ4 | CQHRSTWVTF | 8 |
| S8 | S8V2-112 | IGHV4-59 | IGHD3-10 | IGHJ6 | CARVSMVQG**EGW**YYGMDVW | 17 | N.R. | N.R. | N.R. | N.R. |
| S8 | S8V2-113 | IGHV4-59 | IGHD3-10 | IGHJ6 | CARVSMVLG**EGW**FHGMDVW | 17 | IGLV3-1 | IGLJ3 | CQAWDSNTVVF | 9 |
| S8 | S8V2-117 | IGHV4-59 or IGHV4-61 | IGHD3-10 | IGHJ6 | CARISMVLG**EGW**YHGMDVW | 17 | N.R. | N.R. | N.R. | N.R. |

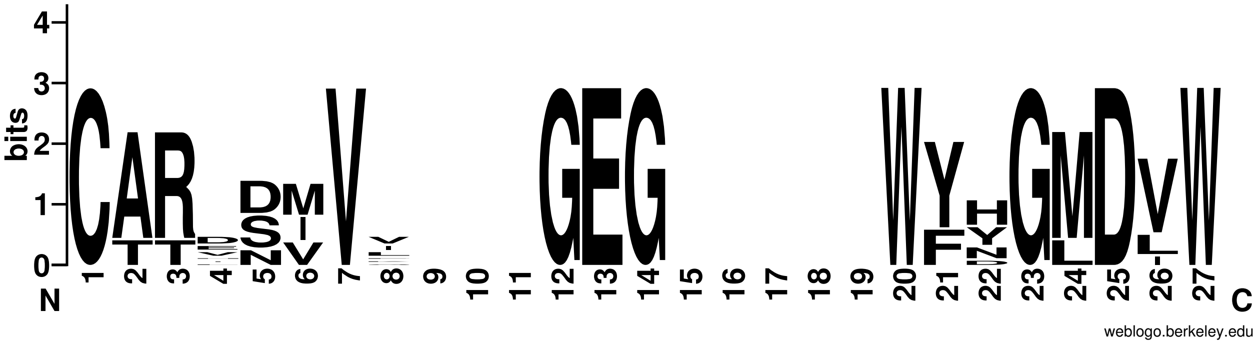


B

| Donor | Antibody | IGHV | IGHD | IGHJ | HCDR3 Sequence | HCDR3 length | IgLV | IGLJ | LCDR3 | LCDR3 Length |
|---|---|---|---|---|---|---|---|---|---|---|
| S5 | S5V2-28 | IGHV3-15 | IGHD2-15 | IGHJ3 or IGHJ6 | CATENVVVG**E**H**W**YRGFDVW | 17 | IGLV2-14 | IGLJ2 or IGLJ3 | CSSYTNTNTLDVVF | 12 |
| S5 | S5V2-34 | IGHV3-15 | IGHD2-15 | IGHJ6 | CTTENVVEG**E**H**W**YHGLDVW | 17 | IGLV2-14 | IGLJ2 or IGLJ3 | CSSYAXTNTLXVVF | 12 |
| S5 | S5V2-35 | IGHV3-15 | IGHD2-15 | IGHJ6 | CTTENVVVG**E**S**W**YYGLDVW | 17 | IGLV2-14 | IGLJ2 or IGLJ3 | CSSYTSISTLDVVF | 12 |
| S5 | S5V2-44 | IGHV3-15 | IGHD2-15 | IGHJ6 | CTTENVVVG**E**H**W**YHGLDVW | 17 | IGLV2-14 | IGLJ2 or IGLJ3 | CSSYTNMNTLDVVF | 12 |
| S5 | S5V2-46 | IGHV3-15 | IGHD2-15 | IGHJ6 | CCTENVVVG**E**N**W**YYGLDVW | 17 | IGLV2-14 | IGLJ2 or IGLJ3 | CCSYSSIITLDVVF | 12 |
| S5 | S5V2-51 | IGHV3-15 | IGHD2-15 | IGHJ6 | CTTENVVVG**E**N**W**YHGLDVW | 17 | IGLV2-14 | IGLJ2 or IGLJ3 | CSSYTNMNTLDVVF | 12 |
| S5 | S5V2-67 | IGHV3-48 | IGHD4-17 | IGHJ6 | CARFDEDA**EG**YYGMDVW | 15 | IGKV1-5 | IGKJ1 | CQQYNAYWTF | 8 |
| S5 | S5V2-73 | IGHV4-61 | IGHD5-18 | IGHJ6 | CARDMLHFDRYSYGYSPDYYYYGMDVW | 25 | IGLV2-14 | IGLJ2 or IGLJ3 | CSSYTNINTLDVVF | 12 |
| S5 | S5V2-74 | IGHV3-15 | IGHD2-15 | IGHJ6 | CTTEDVVVG**E**N**W**YNGMDVW | 17 | IGLV2-14 | IGLJ2 or IGLJ3 | CSSYTSSSTLDVVF | 12 |
| S5 | S5V2-77 | IGHV3-30 | IGHD2-21 | IGHJ4 | CAKAGNAGWECGSDCYEGDYW | 19 | IGLV2-14 | IGLJ1 | CSSHASSSPYVF | 10 |
| S5 | S5V2-81 | IGHV3-48 | IGHD4-17 | IGHJ6 | CARFDEDA**EG**YYGMDVW | 15 | IGKV1-5 | IGKJ1 | CQQYNFYWTF | 8 |
| S5 | S5V2-82 | IGHV3-15 | IGHD2-15 | IGHJ5 or IGHJ6 | CITENVVVG**E**H**W**YNGFDVW | 17 | IGLV2-14 | IGLJ2 or IGLJ3 | CSAYTTTNTLDVVF | 12 |
| S5 | S5V2-86 | IGHV3-15 | IGHD2-15 | IGHJ6 | CTTENVVVG**E**H**W**YHGMDVW | 17 | IGLV2-14 | IGLJ2 or IGLJ3 | CSSYTNINTLDVVF | 12 |
| S5 | S5V2-98 | IGHV3-15 | IGHD2-15 | IGHJ6 | CTTENVVVG**E**H**W**FYGLDIW | 17 | IGLV2-14 | IGLJ2 or IGLJ3 | CSSYTTSGTLDVVF | 12 |
| S5 | S5V2-99 | IGHV3-30 | IGHD3-16 | IGHJ6 | CARADEDV**EG**YYGMDVW | 15 | IGKV1-5 | IGKJ2 | CQQYNNYPHTF | 9 |
| S5 | S5V2-100 | IGHV3-15 | IGHD2-15 | IGHJ6 | CTTENVVVG**E**H**W**YHGLDVW | 17 | IGLV2-14 | IGLJ2 or IGLJ3 | CSSYTTATTLDVVF | 12 |
| S5 | S5V2-103 | IGHV3-15 | IGHD2-15 | IGHJ6 | CTTENVVVG**E**H**W**YHGLDVW | 17 | IGLV2-14 | IGLJ2 or IGLJ3 | CSSYTTATTLDVVF | 12 |
| S5 | S5V2-107 | IGHV3-15 | IGHD2-15 | IGHJ6 | CTTENVVVG**E**H**W**FNGMDVW | 17 | IGLV2-14 | IGLJ2 or IGLJ3 | CSSYTNINTLDVIF | 12 |
| S5 | S5V2-112 | IGHV3-48 | IGHD5-12 | IGHJ6 | CATSRPSGYDG**EG**FFYYYYGMDVW | 22 | IGKV1-9 | IGKJ3 | CQQLNSPFTF | 8 |
| S5 | S5V2-113 | IGHV3-48 | IGHD4-17 | IGHJ3 or IGHJ6 | CARFDEDA**EG**HYGMDVW | 15 | IGKV1-5 | IGKJ1 | CQQYNVYWTF | 8 |
| S5 | S5V2-116 | IGHV3-48 | IGHD4-17 | IGHJ6 | CARFDEDA**EG**YYGMDVW | 15 | IGKV1-5 | IGKJ1 | CQQYNAYWTF | 8 |
| S8 | S8V1-126 | IGHV1-69 or IGHV1-69D | IGHD3-10 | IGHJ5 | CATTHPPRG**EG**VVLGIIEKGFDPW | 22 | IGKV1-39 or IGKV1D-39 | IGLJ2 or IGLJ3 | CQQTFGAPYNF | 9 |
| S8 | S8V1-140 | IGHV3-33 | IGHD5-12 | IGHJ6 | CARTDMDV**EG**YYGMDVW | 15 | IGKV1-5 | IGKJ1 | CQQYYTYSWTF | 9 |
| S8 | S8V1-170 | IGHV3-33 | IGHD5-12 | IGHJ6 | CARTDMDV**EG**YYGMDVW | 15 | IGKV1-5 | IGKJ1 | CQQYYTYSWTF | 9 |
| S8 | S8V2-14 | IGHV1-69 or IGHV1-69D | IGHD3-3 | IGHJ5 | CARSHPPGG**EG**VVFGILEKGFDPW | 22 | IGKV1-39 or IGKV1D-39 | IGKJ2 | CQQTYGSPYTF | 9 |
| S8 | S8V2-56 | IGHV3-13 | IGHD3-9 | IGHJ6 | CAGGDYDI**EG**YYGMDVW | 15 | IGKV1-5 | IGKJ4 | CQQYYSYPLTF | 9 |
| S8 | S8V2-58 | IGHV3-23 | IGHD6-19 | IGHJ4 | CAKEDIAVA**G**YYFDYW | 14 | IGKV3-11 | IGKJ2 | CQQRSNWPRCSF | 10 |
| S8 | S8V2-88 | IGHV3-33 | IGHD2-2 | IGHJ6 | CARIDMDV**EG**YYAMDVW | 15 | IGKV1-5 | IGKJ1 | CQHYDHYSWTF | 9 |
| S8 | S8V2-124 | IGHV3-33 | IGHD2-2 | IGHJ6 | CGRLDMDV**EG**YYGMDVW | 15 | IGKV1-5 | IGKJ1 | CQHYDHYSWTF | 9 |

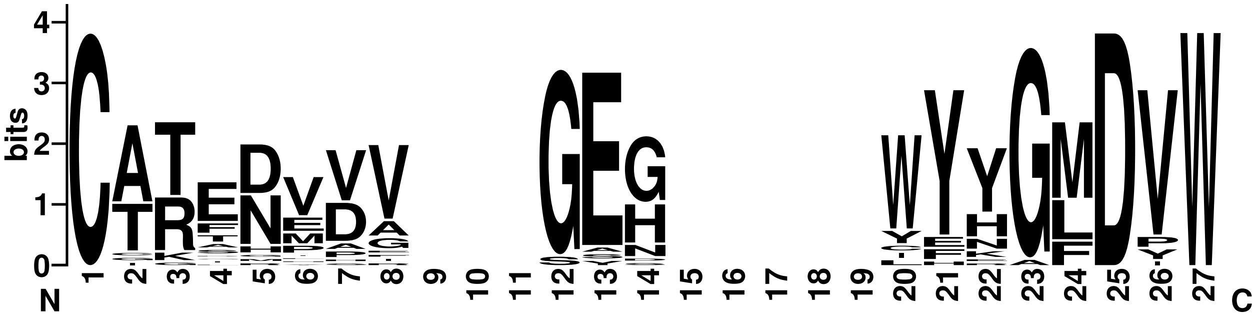

### Supporting Figure 3

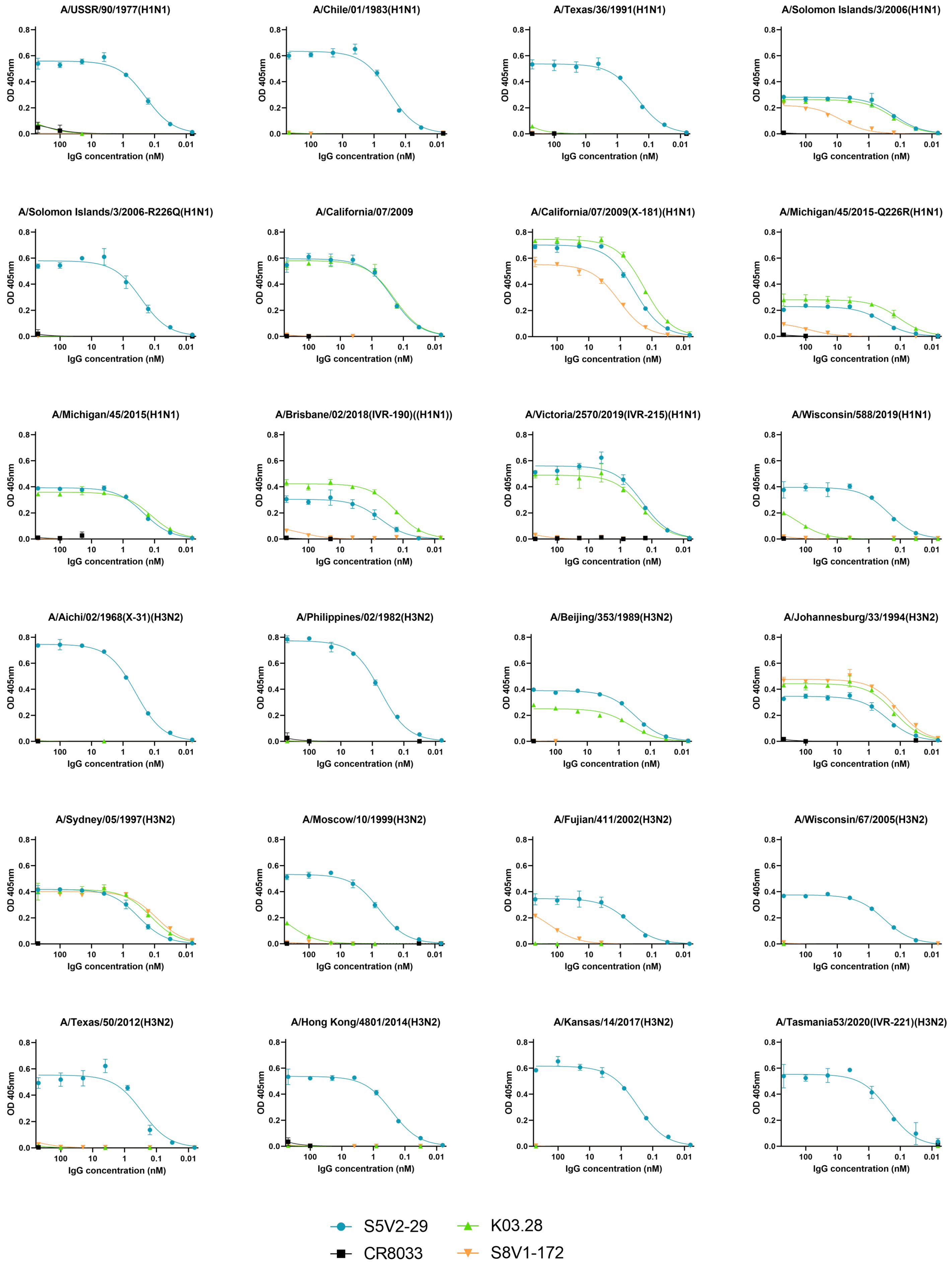

### Supporting Figure 4

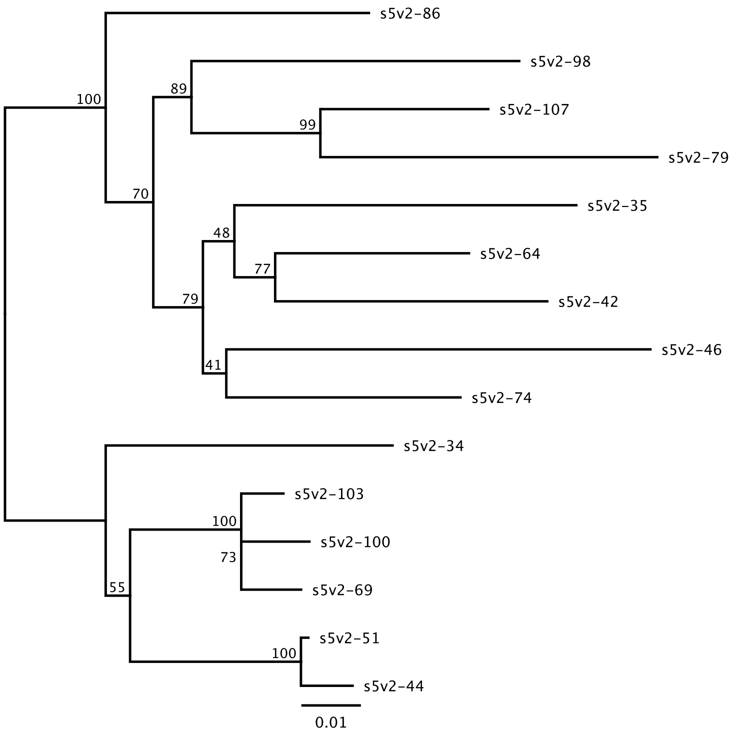


UCA:     HCDR3:CTTENVVVGEHWYYGMDVW  LCDR3:CSSYTSSSTLDVVF
S5V2-86: HCDR3:CTTENVVVGEHWYHGMDVW  LCDR3:CSSYTNINTLDVVF
S5V2-98: HCDR3:CTTENVVVGEHWFYGLDIW  LCDR3:CSSYTTSGTLDVVF
S5V2-107:HCDR3:CTTENVVVGEHWFNGMDVW  LCDR3:CSSYTNINTLDVIF
S5V2-79: HCDR3:CTTENVVVGEGWFNGMDVW  LCDR3:CSSYTNTNTLDVVF
S5V2-35: HCDR3:CTTENVVVGESWYYGLDVW  LCDR3:CSSYTSISTLDVVF
S5V2-64: HCDR3:CATENVVVGEHWYYGMDVW  LCDR3:CSSYTSITTLDVVF
S5V2-42: HCDR3:CTTENVVVGEGWYYGMDVW  LCDR3:CSSYGSITTLDVVF
S5V2-46: HCDR3:CCTENVVVGENWYYGLDVW  LCDR3:CCSYSSIITLDVVF
S5V2-74: HCDR3:CTTEDVVVGENWYNGMDVW  LCDR3:CSSYTSSSTLDVVF
S5V2-34: HCDR3:CTTENVVEGEHWYHGLDVW  LCDR3:CSSYAXTNTLXVVF
S5V2-103:HCDR3:CTTENVVVGEHWYHGLDVW  LCDR3:CSSYTTATTLDVVF
S5V2-100:HCDR3:CTTENVVVGEHWYHGLDVW  LCDR3:CSSYTTATTLDVVF
S5V2-69: HCDR3:CTTENVVVGEHWYHGLDVW  LCDR3:CSSYTTATTLDVVF
S5V2-51: HCDR3:CTTENVVVGENWYHGLDVW  LCDR3:CSSYTNMNTLDVVF
S5V2-44: HCDR3:CTTENVVVGEHWYHGLDVW  LCDR3:CSSYTNMNTLDVVF

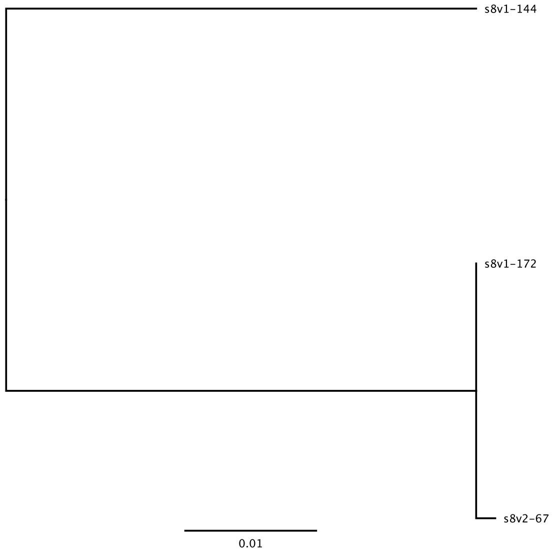


UCA-1:    HCDR3: CARDDIVIGEGWYHGMDLW  LCDR3:CQHRSTWVTF
UCA-2:    HCDR3: CARDDIVIGEGWYHGMDLW  LCDR3:CQHGSTWPTF
S8V1-144: HCDR3: CARDDIVVGEGWYNGMDVW  LCDR3:CQHGSTWPTF
S8V1-172: HCDR3: CARDDIVIGEGWYHGMDLW  LCDR3:CQHRSTWVTF
S8V2-67:  HCDR3: CARDDIVIGEGWYHGMDLW  LCDR3:CQHRSTWVTF

### Supporting Figure 5

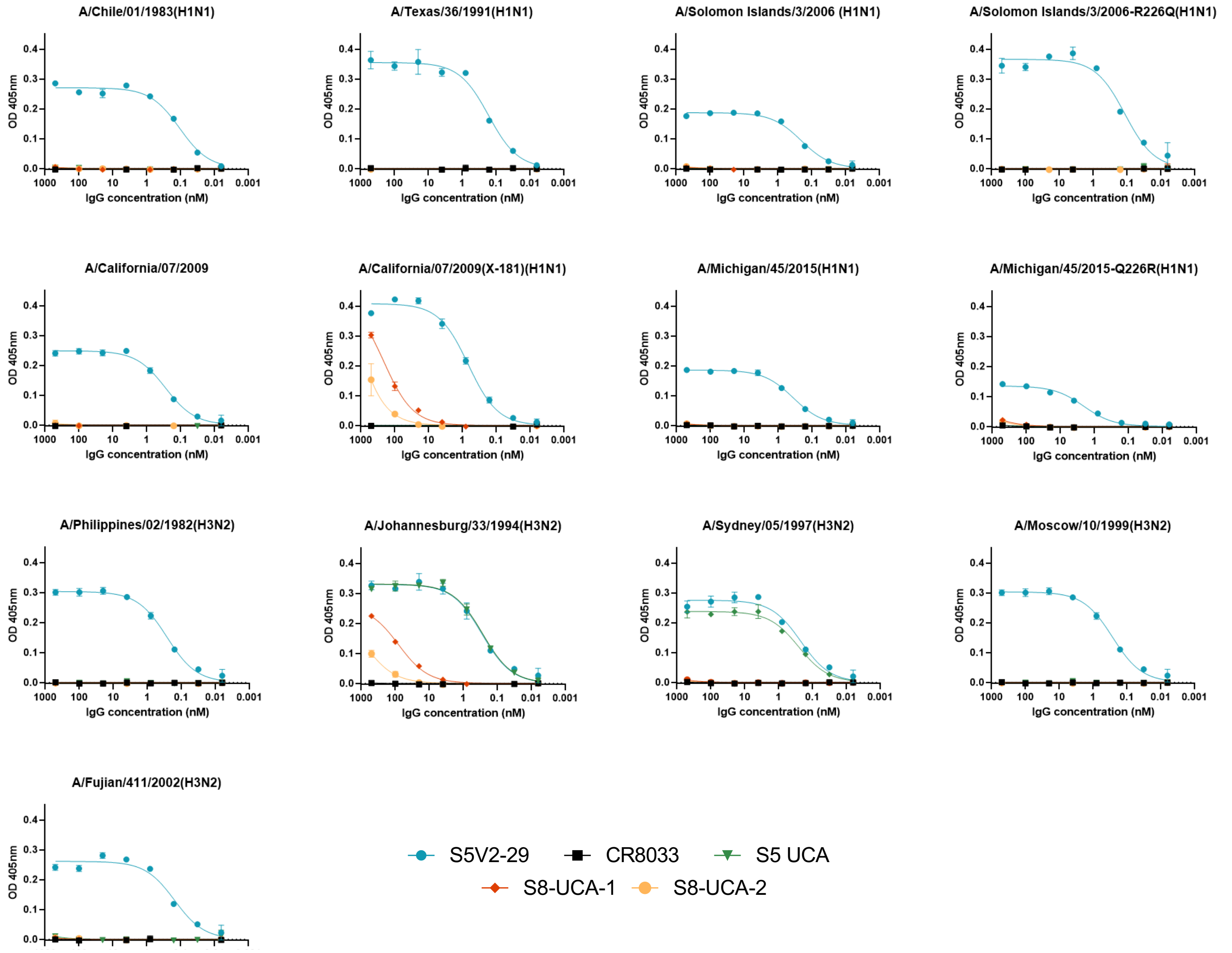


| | | | Antibody | | | | |
|---|---|---|---|---|---|---|---|
| | | Pos. 226 | S8-UCA-1 | S8- UCA-2 | S5 UCA | CR3022 | S5V2-29 |
| H1N1 | A/Chile/01/1983 | Q | >100 | >100 | >100 | >100 | 0.11 |
| | A/Texas/36/1991 | Q | >100 | >100 | >100 | >100 | 0.17 |
| | A/Solomon Islands/3/2006 | R | >100 | >100 | >100 | >100 | 0.20 |
| | A/Solomon Islands/3/2006-R226Q | Q | >100 | >100 | >100 | >100 | 0.12 |
| | A/California/04/2009 | Q | >100 | >100 | >100 | >100 | 0.28 |
| | A/California/07/2009(X-181) | R | >100 | >100 | >100 | >100 | 0.68 |
| | A/Michigan/45/2015 | Q | >100 | >100 | >100 | >100 | 0.36 |
| | A/Michigan/45/2015-Q226R | R | >100 | >100 | >100 | >100 | 2.1 |
| H3N2 | A/Philippines/02/1982 | L | >100 | >100 | >100 | >100 | 0.27 |
| | A/Johannesburg/33/1994 | Q | 81 | >100 | 0.27 | >100 | 0.29 |
| | A/Sydney/05/1997 | I | >100 | >100 | 0.26 | >100 | 0.22 |
| | A/Moscow/10/1999 | V | >100 | >100 | >100 | >100 | 0.27 |
| | A/Fujian/411/2002 | V | >100 | >100 | >100 | >100 | 0.15 |

| Kd in nM | | | | |
|---|---|---|---|---|
| <1 | 1-5 | 5-25 | 25-100 | >100 |

### Supporting Table 1

| | K03.28-A/California/7/2009 (H1N1) (NYMC-X181) (H1-X181) Fab-HA head complex | S8V1-172-A/Sydney/05/1997(H3N2) Fab-HA head complex |
|---|---|---|
| **PDB ID** | 7TRH | 7TRI |
| **Data Collection** | APS 24-ID-C | APS 24-ID-C |
| Number of datasets | 1 | 1 |
| Resolution, Å | 46.80-3.00 (3.11-3.0) | 47.35-3.60 (3.73-3.60) |
| Wavelength (Å) | 0.9791 | 0.9791 |
| Space Group | P 1 21 1 | P 32 2 1 |
| Unit cell dimensions (a, b, c), Å | 75.83 43.65 96.91 | 94.7 94.7 248.84 |
| Unit cell angles (α, β, γ) ° | 90.00 105.03 90.00 | 90.00 90.00 120.00 |
| I/σ | 14.93 (2.10) | 12.72 (3.04) |
| Rmeas | 0.06 (0.54) | 0.06 (0.46) |
| Rpim | 0.04 (0.38) | 0.046 (0.32) |
| Rmerge, % | 0.04 (0.38) | 0.045 (0.32) |
| CC* | 1.00 (0.93) | 1.00 (0.91) |
| CC½ | 1.00 (0.77) | 1.00 (0.71) |
| Completeness, % | 98.33 (97.87) | 99.46 (99.87) |
| Number of reflections | 24401 (2401) | 30957 (3053) |
| Redundancy | 2.0 (1.9) | 2.0 (2.0) |
| **Refinement** | | |
| Number of reflections: | | |
| Working | 12440 (1241) | 15605 (1543) |
| Free | 623 (62) | 789 (86) |
| Rwork, % | 20.20 (31.45) | 23.73 (31.01) |
| Rfree, % | 24.32 (35.44) | 26.66 (34.98) |
| Ramachandran plot, % (favored, disallowed) | 93.42 (0.16) | 93.42 (0.14) |
| Rmsd bond lengths, Å | 0.003 | 0.003 |
| Rmsd bond angles, ° | 0.64 | 0.61 |
| Average B-factor | 78.52 | 119.17 |

Rmerge, Σh,k,l|Σi|Ii(hkl) – <I(hkl)>|/Σh,k,lΣiIi(hkl), where I is an intensity that is observed i times; I/σ, signal-to-noise ratio (average observed intensity divided by average standard deviation of the observed intensity); Rwork, Σh,k,l||Fobs| – |Fcalc||/Σh,k,l|Fobs|, where h, k, l covers the "working set" of observed structure factor amplitude (Fobs) reflections used in refinement (total reflections minus the test set) and Fcalc is the calculated structure factor amplitude; Rfree, calculated as for Rwork but on 5% of data excluded prior to refinement. Values in parentheses refer to highest-resolution shell. Related to Experimental Procedures.
